# Decomposition potential and population structure among active microbial communities in deep salt marsh sediments

**DOI:** 10.1101/2025.07.18.665577

**Authors:** Joseph H. Vineis, Ashley N. Bulseco, Sa’ad A. A. Rafie, Zoe G. Cardon, Jennifer L. Bowen

## Abstract

Salt marsh sediments contain high levels of microbial diversity and function that reflect the heterogeneity of available nutrients, dynamic hydrology, and layers of organic matter developed over millennia. However, the microbially mediated processes involved in the cycling of complex carbon in these sediments are still largely undescribed. We used genome reconstruction, cooccurrence networks, and genome scale metabolic modeling to identify the functional capacity for organic matter decomposition among cooccurring microbes from 240 cm of sediment in a *Spartina patens* dominated salt marsh, representing over 3000 years of sediment accumulation. Microbial communities were structured according to depth, and the network identified significant positive cooccurrence within the community. We identified an active subnetwork of cooccurring subsurface metagenomic assembled genomes (MAGs) capable of decomposing complex carbon and aromatics. The Wood-Ljungdahl pathway was central to most of the MAGs within the subnetwork and metabolism of aromatics was potentially enabled through metabolic handoffs and the Benzoyl-CoA pathway. Variant analysis indicates that these subsurface populations are stratified with depth, suggesting high selection coupled with spatial isolation. These results provide evidence of microbial populations that persist under energy limited conditions and contribute to the transformation of organic matter within salt marsh sediment.

## Introduction

Salt marshes are globally important to carbon storage and have the potential to influence both the changing climate and the stability of coastal systems. Salt marsh sediments contain between 400 – 6500 Tg of organic carbon (Duarte et al. 2013; Mcleod et al. 2011) and store an additional 10.2 - 44.6 Mt annually, the equivalent of 0.5-1.0% of anthropogenic C emissions (Ouyang and Lee 2014). However, carbon storage in salt marshes is declining and estimates indicate that over a recent nine-year period, salt marsh habitat loss resulted in emission of 16.3 Tg of CO_2_ and a reduction of 0.045 Tg of CO_2_ burial per year (Campbell et al. 2022). Rapidly increasing sea level rise and warming climate could destabilize current stocks of organic carbon with unpredictable consequences (Fontaine et al. 2007; Spivak et al. 2019).

Microbial communities within salt marsh sediments are compositionally and metabolically diverse and their interactions influence pools of soil organic carbon (SOC) (Bulseco et al. 2019; Erb et al. 2024). However, there are important knowledge gaps in our understanding of the metabolic capacity, distribution, and processes that shape microbial communities and populations in blue carbon systems. Genomic characterization of microbial communities can help to fill these gaps and often leads to the identification of previously unidentified carbon cycling among novel taxa (Payne et al. 2025; Seitz et al. 2016; Vineis et al. 2023). Identifying how pools of carbon are influenced by microbial communities will enhance our understanding of the fate of blue carbon stocks (Hilmi et al. 2021), how they are formed, and how they change over time.

The decomposition of organic material is strongly influenced by plant communities that blanket the surface of the marsh in nearly homogenous stands. The regions of the marsh inhabited by plants are highly dynamic zones that contain hot spots of microbial activity due to plant-microbial interactions and feedbacks that are sensitive to seasonal and tidal influence (Kuzyakov and Blagodatskaya 2015). *Spartina patens* is a highly productive salt marsh macrophyte with roots that release organic carbon, thereby influencing carbon sequestration of salt marshes through the “microbial carbon pump” (Liang et al. 2017; Spivak et al. 2019). Live roots of *S. patens* are dominantly 0-10 cm below the sediment surface (Gallagher and Plumley 1979) yet plant inputs could influence the SOC pool down to 100 cm (Liu et al. 2017). Additionally, the tidal cycle causes redox gradients to shift rapidly, and oxygenated water can infiltrate deeper sediments due to bioturbation and radial oxygen loss from roots (Haviland and Noyce 2024), resulting in a dynamic distribution of resources within the zone of tidal and plant influence. These dynamic conditions shape the microbial strategies for respiration and decomposition.

The patchy distribution of resources and dynamic processes in marsh sediments present challenges for unraveling the underlying rules for community assembly and metabolism. Identifying the distribution and putative functions of the microbial genomes within the dynamic zone influenced by plants and tidal inputs remains one of the most challenging but critical aspects to understanding the carbon and nutrient cycling in salt marsh systems. While decomposition within the salt marsh is conducted primarily by sulfate reducers (Howes et al. 1984), most of the alternative pathways for anaerobic respiration, fermentation, and carbon fixation have also been identified in marsh sediments (Baker et al. 2015; Bulseco et al. 2020; Vineis et al. 2023). We are only beginning to understand how these metabolic processes are linked among organisms and within individual genomes.

Below the rooting zone and beyond the influence of living plants, salt marshes have accumulated vast stores of organic peat that date back, in some cases, millennia (Kirwan et al. 2023). These older sediments are likely to contain microbes adapted to survive in deep, low energy environments where prior oxidation of organic material has led to a decline in the bioavailability of organic carbon with sediment age (Arndt et al. 2013). Microbial communities below the influence of living plants therefore might mirror other marine low energy sediment communities supported by complex organic matter derived from compounds such as lignin, cellulose, and phenolic polymers (Jørgensen and Boetius 2007). Decomposition of these compounds within energy limited sediment is often dependent on syntrophic interactions or metabolic handoffs among microbial consortia (Peng et al. 2023). The specific mechanism of microbial decomposition has important consequences for carbon stability because microbial turnover can result in the production of carbon that is protected from further decomposition (Liang et al. 2017). Therefore, identifying the metabolic potential and activity of microbial decomposition is important to understanding the long-term stability of organic carbon in salt marsh sediment.

Due to the gradient of bioavailable carbon in salt marsh sediments, we hypothesize that 1) microbial communities will be structured according to sediment depth, a proxy for sediment age; 2) deeper communities will be functionally streamlined due to environmental filtering; 3) the community found within deeper sediments will be more likely to contain cooccurring members; 4) there will be metabolic complimentary enabling sequential decomposition of complex carbon among microbial communities located in deeper sediments; 5) the population structure will be depth-dependent, similar to patterns observed in other energetically limited marine environments (Kirkpatrick et al. 2019; Walsh et al. 2016).

## Methods

We employed a stepwise methodological approach that integrates genome reconstruction, cooccurrence networks, genome scale metabolic models, and metabolic complementarity to enable the identification of cooperative potential for carbon cycling microbial consortia (Fig. 1). We began with microbial genome reconstruction that relied heavily on manual curation of each metagenome assembled genome (MAG). A manual approach is rarely conducted at this scale because automated approaches are far faster and less labor intensive, but visual inspection is often required to reduce the recruitment of incorrect sequence information into MAGs. Additionally, MAG reconstruction is often conducted on a co-assembly of multiple samples, but we used individual assemblies to avoid the potential of creating chimeric contigs and to evaluate the presence of novel genomic features for future work. This step also allows for dereplication to identify populations that occur across multiple samples, complementing the often-employed read mapping used to assess MAG relative abundance. In the second step, we generated cooccurrence networks to identify significantly associated cooccurring microbes. Cooccurrence in our analysis does not imply a direct spatial or physical relationship and additional evidence beyond genomic analysis is required to identify spatial connections. This analysis led to the identification of network connectors, MAGS that were positively connected to abundant members of cooccurring network modules primarily found below 40 cm. We then used a metabolic complementarity analysis and metatranscriptomics to address the likelihood of metabolic handoffs within this cooccurring subnetwork. Finally, we applied a single nucleotide variant analysis of a subset of genomes within the subnetwork to identify underlying structure at the population level. We outline the details of all steps in our approach below and how they are used to test each hypothesis. Additional details of our analysis can be found in the git associated with this manuscript (https://github.com/jvineis/DEEP-CORE-MANUSCRIPT.git).

**Figure 1.**
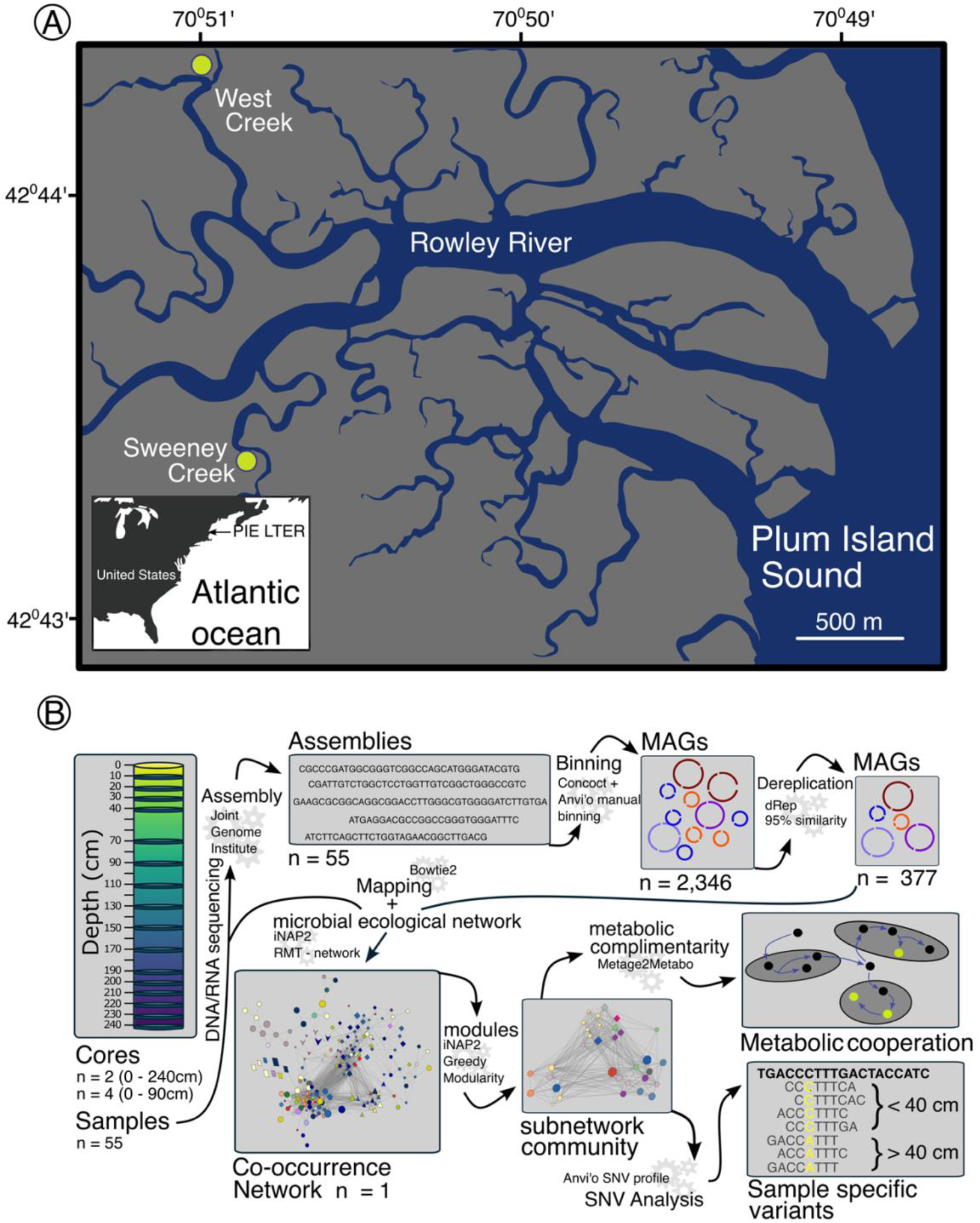
A) Map of the coring locations within the Plum Island Estuary Long Term Ecological Research (PIE LTER) site and B) Details of core sampling and the bioinformatic approaches used in this study. The arrows flow from the sampling approach through genome reconstruction, network analysis, identification of a cooccurring subnetwork, and the metabolic complementarity and variant detection of that subnetwork. The primary bioinformatic tool is shown for each step.

### 1. Sampling

Six sediment cores were taken using a Russian peat corer from the salt marsh platform at two sites in Rowley MA, USA (42.759 N, 70.891 W) primarily vegetated by *Spartina patens* in July of 2016. The sites are part of the Plum Island Ecosystem (PIE) Long-Term Ecological Research site (LTER). Three cores were collected from the high marsh *Spartina patens* platform at each of two creeks, West and Sweeney Creek (Fig. 1A). The two creeks are approximately 2 km apart and cores collected within each creek were separated by less than 50 meters. One core at each site was taken to the point of refusal, reaching a maximum depth of 240 cm, and the other two cores at each site were taken to approximately 100 cm depth. Once extracted, subsections of the core were collected approximately every 10 cm and a 2 cm sub-section was homogenized in a sterile 50 mL falcon tube, flash frozen using liquid nitrogen and then stored at -80°C for nucleic acid analysis. A detailed description of sample handling and analysis, including biogeochemistry of the cores, has been previously described (Bulseco 2018).

### 2. Metagenomic and Metatranscriptomic Sequencing

The input DNA for sequencing library construction was extracted from homogenized sediment from each 2 cm thick fraction (0.25 g wet weight) using a Qiagen Power Soil DNA extraction kit (Qiagen, Germantown, MD) according to the manufacturer’s recommendations. We evaluated the quality and the quantity of DNA using a Nanodrop and Quant-IT DNA assay (Invitrogen, USA) respectively. RNA was extracted using the Qiagen RNeasy Power Soil Kit (Qiagen, Germantown, MD). The quantity was evaluated using the Quant-IT RNA assay (Invitrogen, USA) and quality evaluated on the TapeStation (Agilent, USA). Purified DNA and RNA was shipped overnight to the Joint Genome Institute (JGI) on dry ice. At JGI, DNA and RNA libraries were constructed according to standard methods. Each library was uniquely barcoded and sequenced according to PE 2×151 chemistry on an Illumina NovaSeq. Sequences were quality filtered using the default settings of BBTools v38.26 at JGI (Bushnell 2014). Quality filtered DNA sequences from each metagenomic sample were independently assembled using SPAdes assembler 3.12.0 (Bankevich et al. 2012).

### 3. MAG reconstruction, dereplication, phylogenomics and metagenomic read recruitment

We used DNA assemblies and quality filtered sequences generated by JGI/IMG for each of the 55 samples to reconstruct MAGs for each independent sample (Fig. 1B). MAG reconstruction efforts can be improved by co-assembly of multiple samples in a gradient or time series, especially when read counts are not sufficient to provide the minimum amount of coverage necessary for assembly (Stewart et al. 2018). However, the detection of redundant MAGs across samples requires that genome reconstruction be carried out for each sample independently. Additionally, the number of high-quality genomes recovered can be improved by reconstructing genomes from individual samples compared to co-assembly of multiple samples (Olm et al. 2017).

We reconstructed draft genomes for each sample using CONCOCT (Alneberg et al. 2014) as an automated guide for genome binning, followed by manual curation of each CONCOCT MAG in Anvi’o, using contig tetranucleotide frequency, coverage, taxonomic assignment, completion, and contamination to guide this process (Eren et al. 2015, 2020). In cases where the placement of contigs could not be resolved among a group of similar MAGs, we used anvi-refine to provide increased resolution on this subset of similar MAGs. This allowed for the separation of closely related MAGs and/or the improvement of draft genome bins, as determined by Anvi’o real-time completion and contamination scores. An example of the manual binning process and MAG refinement can be found in the git associated with this manuscript https://github.com/jvineis/DEEP-CORE-MANUSCRIPT.git.

Dereplication can indicate population structure because reconstruction of the same MAG in multiple samples is indicative of its distribution within the environment. Therefore, the level of dereplication is helpful to address hypothesis 1 (communities are structured by depth) and hypothesis 5 (populations are structured with depth). Dereplication of the MAGs and selection of the representative MAG from the dereplicated collection were conducted using the default settings for dRep v 2.0.0 (Olm et al. 2017). Default settings in dRep include dereplication of genomes with an average nucleotide identity (ANI) of 0.95 and a minimum overlap of 10% between the two genomes. dRep incorporates additional essential methods, including CheckM v1.0.7 (Parks et al. 2015) and FastANI (Jain et al. 2018). dRep uses completion and contamination based on checkM. MAG completion threshold was never at or above the 75% for any of the Patescibacteria, which were dereplicated separately without a checkM quality score filter. Any MAG with a completion score below 50% and contamination above 10% according to Anvi’o estimates were removed from downstream analysis. We used the GTDB-Tk v 2.1.1 to assign taxonomy to each of the MAGs recovered using default settings (Chaumeil et al. 2020).

To identify the phylogenomic relationships among dereplicated MAGs and the presence of novel taxa, we first exported the amino acid sequence of the best hit for 71 single copy genes (Bacteria_71) (Lee 2019), aligned them using MUSCLE and trimmed them using trimAl v1.4 (Capella-Gutiérrez et al. 2009). We reconstructed the phylogenetic tree using the concatenated sequence alignment according to anvi-gen-phylogenomic-tree which employs the FastTree tree building algorithm (Price et al. 2009). The fasta file of single copy gene sequences, alignment, tree, and additional details of phylogenetic reconstruction are available at github https://github.com/jvineis/DEEP-CORE-MANUSCRIPT.git.

We mapped the metagenomic reads from each sample to the collection of dereplicated MAGs using bowtie2 v2.5.3 (Langmead and Salzberg 2012). Unmapped reads were filtered from the results and samtools v1.19.2 (Li et al. 2009) was used to tabulate the number of reads recruited to each MAG. The relative abundance of each MAG was estimated by dividing the number of reads mapped to the MAG by the number of reads recruited to the non-redundant collection of MAGs. We did not use the total reads sequenced as the denominator because of the potential for eukaryotic contamination in these samples, especially in shallow sediments (Sullivan and Currin 2002; Kearns et al. 2019). To estimate the distribution of each MAG across the depth profile, we weighted the relative abundance observed for each MAG by the number of samples collected at that depth (depth weighted abundance). Because we were not interested in comparing the relative abundance of MAGs within samples, no normalization to correct for genome size was employed. The proportion of relative abundance was calculated for each depth by summing the total relative abundance for a MAG and dividing it by the sum of depth weighted abundance. Although we did not generate a specific statistic to test for proportional differences of each MAG with depth, this provides a descriptive guide to the distribution of the MAGs recovered in our study.

### 4. Community beta diversity and Identifying cooccurring MAGs using network reconstruction

We employed a beta diversity analysis based on the read recruitment from metagenomic samples to the collection of dereplicated MAGs. This is a direct test of hypothesis 1 (communities are structured by depth). The count matrix of reads per MAG was used as input to compute a Bray-Curtis dissimilarity matrix followed by non-metric multidimensional scaling. The betadisper function was used to test for differences in dispersion among cores and depths. Adonis2 was used to test for significant differences in community composition due to depth, core and the interaction between core and depth. All beta diversity analysis was conducted using the vegan package (v2.7.1) (Oksanen et al. 2024) in the R statistical package (v4.5.3) (Core Team 2024).

To identify cooccurring MAGs, we constructed a molecular ecological network (MEN) using random matrix theory (RMT) according to the procedures recommended by iNAP (Feng et al. 2022). We employed this approach and the steps below to test hypothesis 3 (cooccurrence is more common in deep sediments). RMT is more commonly used for functional or phylogenetic marker genes but is broadly designed to identify cooccurrence patterns among complex datasets that contain large numbers of variables (species occurrence, microarray data, marker genes) (Gibson et al. 2013). The count matrix of reads recruited to each of the 377 dereplicated MAGs was used as the input for RMT MEN construction and any zero was converted to 1. Following log transformation of the count matrix, an adjacency matrix was calculated according to Pearson correlation coefficients. To separate significant cooccurrence from noise, RMT estimates the nearest neighbor spacing distribution (NNSD) within the entire correlation matrix. A random matrix of NNSD values will follow a Gaussian distribution and a non-random matrix will follow a Poisson distribution (Luo et al. 2007). To identify the threshold where the correlation estimates transition from Gaussian to Poisson, we ran iterations of successively smaller correlations starting at 1 with a step size of 0.01. A Chi-square test was used to evaluate transition from a Poisson to a Gaussian distribution at each step and iNAP created a plot of resulting p-values to evaluate each step. Visual inspection of the cutoff indicated that correlations below 0.92 were more likely due to random noise (Fig. S1). All correlations below this 0.92 cutoff were set to zero and a network was constructed. RMT network properties were calculated for the resulting filtered network, including R-square of the power-law relationship. A network containing few MAGs with large numbers of connections with other MAGs will have connectivity that follows a power-law distribution. If the connectivity doesn’t fit the power-law distribution, this indicates that there are many MAGs with high connectivity to other MAGs (a very dense network). Module separation and identification of module hubs were calculated according to a greedy modularity optimization (Newman 2004). Within-module (Z) and among-module connectivity (P) were used to classify each MAG as a network hub (Z > 2.5 and P > 0.62), module hub (Z > 2.5 and P < 0.62), connector (Z < and P > 0.62), or peripheral (Z < 2.5 and P < 0.62) (Deng et al. 2012). Based on previous ecological interpretation of networks, the peripheral, connector, module hub, and network hub might represent a continuum from specialist to generalist (Olesen et al. 2006). To identify cooccurring genomes, we focused on modules, interlinked subsets of MAGs, and connectors, which could be important to the stability of the different modules (Olesen et al. 2006; Qin et al. 2023).

### 5. Metabolic potential of MAGs within identified network modules

We annotated all nonredundant MAGs using METABOLIC v4.0 for gene homology and characterization of metabolic pathways. A pathway was considered present if 75% of the genes within the pathway were present in the MAG. Glycoside hydrolases (GH) and polysaccharide lyases (PL) were identified based on gene homology in the Carbohydrate Active Enzyme (CAZy) database (Drula et al. 2022) (http://www.cazy.org/). Genes within each MAG were also queried against MEROPS peptidase database (Rawlings et al. 2007).

Because we identified network modules that were located with a peak abundance at different depths, we analyzed differences in genome properties and functional potential across these modules. Genome size, genome percent completion, number of metabolic pathways, CAZymes, and peptidases were compared among the network modules. The number of CAZymes and peptidases were corrected for number of genes in the MAG. Estimated marginal means were calculated from an analysis of variance using the lm function and we evaluated pairwise differences using the estimated marginal means. Analysis was conducted using emmeans (Lenth 2023) in the R statistical package (v4.5.3) (Core Team 2024). Similarly, the mean number of CAZymes and peptidases were evaluated among network modules.

Identification of the proportion of MAGs containing each of the metabolic pathways and functions within each of the network modules is a test of hypothesis 2 (environmental filtering and genome streamlining of MAGs). This analysis also tests hypothesis 4 (metabolic complementarity and sequential decomposition) because it enables the identification of pathways and genome properties unique to subsurface cooccurring MAGs relevant in other energy depleted environments.

### 6. Genome scale metabolic models, activity, and single nucleotide variant (SNV) analysis of a widespread subsurface subnetwork

Visualization of the complete RMT network using Cytoscape (Shannon et al. 2003) led to the identification of four modules with similar taxonomy, functional profiles, and high relative abundance within deep sediment layers. Bathyarchaeia BA1 MAGs occurred in each of these modules and two of the MAGs were connectors. We identified a subsurface collection of MAGs with significant positive correlations to the Bathyarchaeia BA1 node for further investigation (Bathyarchaeia BA1 subnetwork). Within this subnetwork of associated MAGs, we estimated the metabolic potential for key pathways related to decomposition of organic compounds, aromatics, acetate, hydrogenases, and pathways important to carbon cycling using METABOLIC (v4.0) according to the default parameters (Zhou et al. 2022). METABOLIC-C (v4.0) was used to estimate the metagenomic and metatranscriptomic coverage of functions in this collection of MAGs to calculate the MW-score (metabolic weight score) of present (metagenome) and active (metatranscriptome) functions within each sample. METABOLIC-C was also used to calculate the contribution of each MAG to the cumulative MW-score for each function (Zhou et al. 2022). In the MW-score calculations for the metatranscriptome, mapping data were normalized by gene length and metatranscriptome read count, resulting in reads per kilobase of transcript, per million mapped reads (RPKM). This analysis addresses hypothesis 2 (environmental filtering and genome streamlining of MAGs) and hypothesis 4 (metabolic complementarity and sequential decomposition).

To test for metabolic complementarity among the Bathyarchaeia BA1 subnetwork MAGs, we reconstructed genome-scale metabolic networks (GSMNs) for members of the Bathyarchaeia BA1 subnetwork using gapseq (Zimmermann et al. 2021), and miscoto (Frioux et al. 2018) was employed to screen GSMNs for key members of minimal communities required to produce the metabolites of the entire community. Analysis of GSMNs was employed using Metag2Metabo (M2M) (Belcour et al. 2020). This approach identifies putative metabolites produced by cooccurring members and the key microbial taxa required to produce these metabolites, along with interchangeable (redundant) members of the network. The analysis of genome complementarity is a test of hypothesis 4 (metabolic complementarity and sequential decomposition) and hypothesis 2 (environmental filtering and genome streamlining of MAGs), because it identifies the potential metabolic handoffs important to survival under energy limited conditions. The absence of metabolic handoffs among the community and having only a small number of MAGs be required to produce the metabolites of the entire community would suggest that environmental filtering is the primary driver of cooccurrence.

We investigated sample-specific single nucleotide variant (SNV) profiles for four MAGs from the Bathyarchaeia BA1 subnetwork that had adequate coverage across most samples in the two deepest cores. This analysis tests hypothesis 5 (depth dependent population structure). To identify SNVs within MAGs, we used bam files generated from metagenomic read mapping of each sample to the non-redundant collection of MAGs. SNVs were called according to the heuristic outlined in the anvi-profile command which relies on a minimum departure from the reference sequence to filter low frequency SNVs. To generate the variability profile across samples for each MAG, we used the anvi-gen-variability-profile command in quince mode. We used the anvi-script-snvs-to-interactive to generate a matrix of SNV deviation from the reference for 5000 randomly chosen positions. This matrix of 5000 positions was used to generate Bray-Curtis dissimilarity and test for depth and site dependent differences in SNV frequency. The general relationship between overall variability and depth was plotted for each of the four MAGs according to variability calculation shown in equation 1.

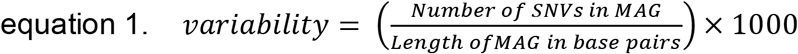

A depth dependent trend in variability could suggest that the population is under high selection where variability is lowest, and the absence of a significant trend indicates similar selection throughout the depth profile and/or unrestricted geneflow (migration through the sediment profile) of the community. A decline in variability with depth and SNVs that are restricted to layers of the sediment is indicative of environmental filtering and restricted gene flow.

## Results

### 1. MAG populations were novel, diverse, and uniquely distributed within the sediment profile

We generated 645.9 Gbp sequences from 55 samples collected from six salt marsh sediment cores (Fig. 1B). The manual genome binning and refinement effort for each of the 55 samples recovered a total of 2,346 MAGs. Dereplication resulted in a collection of 377 MAGs that included 44 phyla (Table S1). The dereplicated MAGs were an average of 87% complete and 2.7% contaminated according to Anvi’o estimates (Fig. 2, Table S1). According to the relative evolutionary distance (RED) score and taxonomic assignment by GTDB-Tk, six MAGs were novel at the order level and included members of the TA06, Proteobacteria, Calditrichota, and Spirochaetia. 5% of MAGs represented novel families and 60% of MAGs received genus level assignment, including 121 unique genera, while less than 1% were assigned at the species level (Table S1).

**Figure 2.**
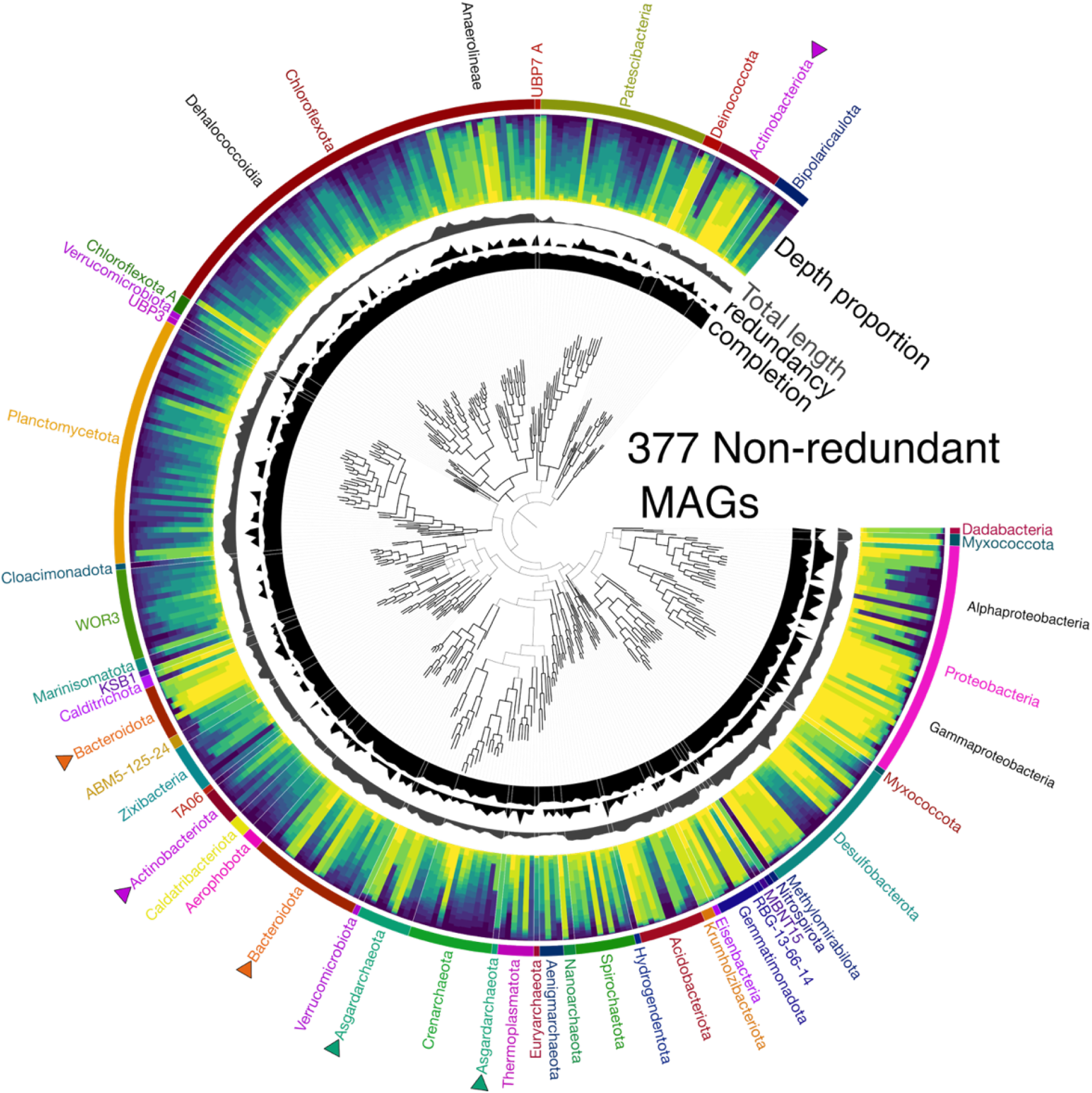
The tree at the center of the display shows the phylogenetic relationship among all MAGs. The innermost ring is a histogram of the single copy gene completion estimates (range = 0-100%), followed by redundancy (range = 0-10%), and total length of the genome (range = 0-7.6Mbp). The “Depth proportion” ring shows the proportion of the total reads mapped to a MAG from each depth. The outermost ring shows the phylum level taxonomy as a color bar with matching text. Arrows are positioned adjacent to phyla names with split positions in the tree.

The proportion of metagenomic sequences recruited (mapped) to our collection of nonredundant MAGs was generally greater in sediment samples collected below 40 cm (Fig. S2). Read recruitment was at least 30% in all but one sample and was over 50% in 16 of the 31 samples below 40 cm. Conversely, in samples collected above 20 cm, less than 20% of sequences were recruited to the nonredundant collection of MAGs (Fig. S2) and the number of MAGs recovered was generally lower. This presents an inherent bias that limits our ability to characterize the more diverse surface sediments, but it allows for a robust subsurface analysis.

The combination of metagenomic read mapping, phylogenetic analysis, and dereplication identified phylogenetic differentiation among MAGs within the sediment. Dominant phyla within sediments below 70 cm included Chloroflexota, Planctomycetota, and WOR3 (Fig. 2). Many MAGs within these three phyla were independently reconstructed from several samples, including 10 Chloroflexota that were reconstructed from at least 5 samples (Fig. S3, Table S1). Within the Chloroflexota, the Anaerolinea were more common in shallow sediments and had larger genome sizes than Dehalococcoidia (Fig. 2). A relationship between genome size and phylogeny was also observed within the Desulfobacterota. WOR-3 MAGs were especially widespread within deeper sediments where we were able to independently reconstruct the same MAG from 30 different samples. Another widespread WOR-3 MAG was independently reconstructed from 26 samples (Fig. S3, Table S1). Nearly all 41 MAGs in the Planctomycetota were more abundant below 40 cm and we identified six that were independently reconstructed from 5 or more samples. (Fig. 2, Fig. S3, Table S1). The 39 Proteobacteria MAGs were split evenly between Alphaproteobacteria, spanning the entire depth gradient, and Gammaproteobacteria, which were most abundant in sediment above 40 cm (Fig. 2). However, within the Proteobacteria we did not observe a strong relationship between genome size and depth and only one Proteobacteria MAG was independently reconstructed from 5 or more samples. The 28 MAGs within the phylum Bacteroidota were partitioned according to depth, although not as strictly as Alpha and Gammaproteobacteria (Fig. 2). This phylum was also split into two separate clades in our phylogenetic analysis. Similarly, separate clades were obtained for Actinobacteriota and Asgardarchaeota (Fig. 2).

### 2. Communities were structured according to depth

We found that communities were significantly structured along the depth gradient (R^2^ = 0.23, p<0.001) and the NMDS ordination displayed a horseshoe like pattern, indicative of an existing environmental gradient (Morton et al. 2017) (Fig 3A, B). We observed a small but significant difference among creeks and the interaction between creek and depth was significant (R^2^ = 0.30) indicating that MAG turnover with depth was different between the two locations (Fig. 3A, B). However, the six cores were generally similar in the overall shift in composition with depth at the phylum level (Fig. 3C).

**Figure 3.**
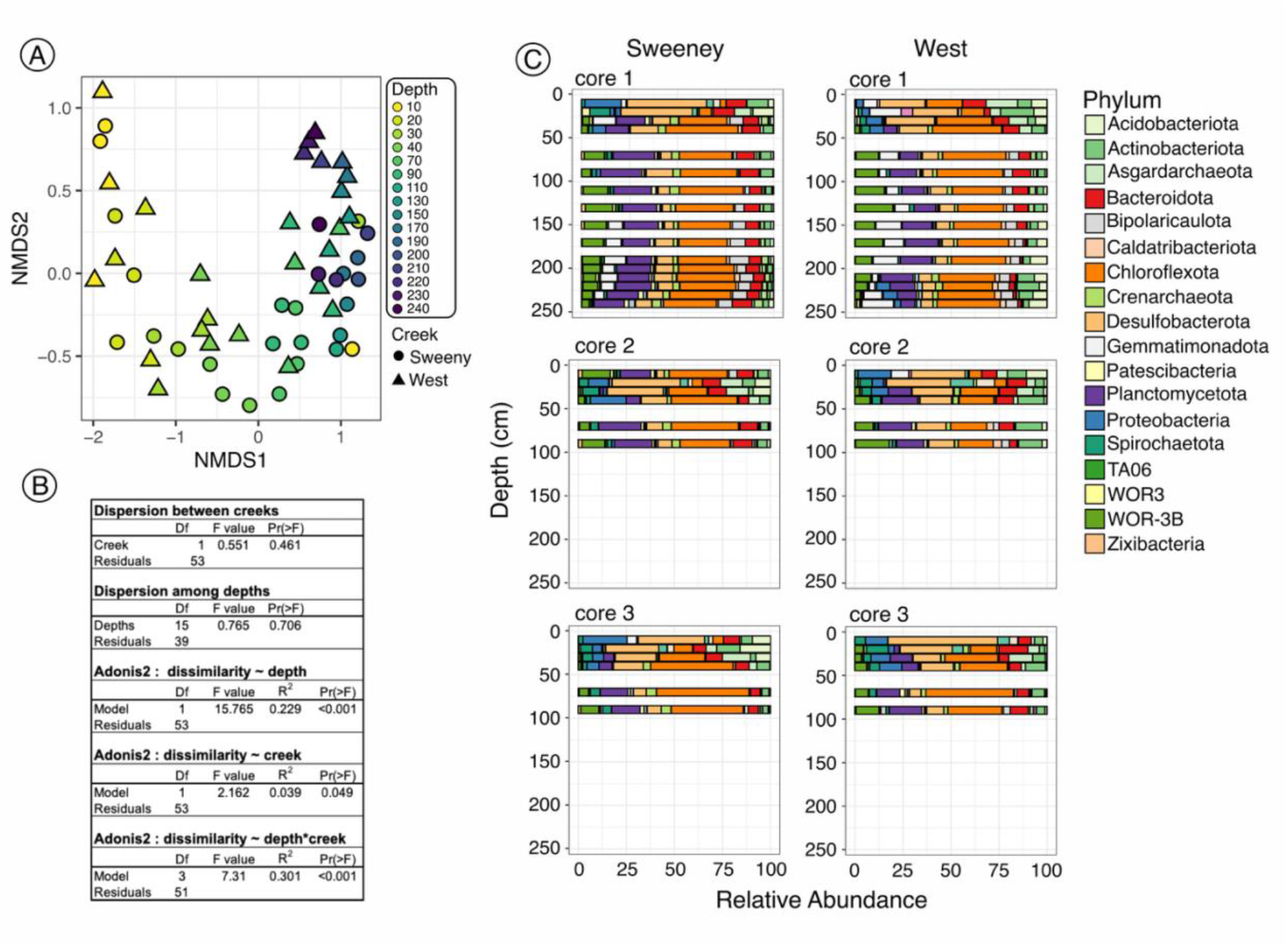
Summary of MAG relative abundance and ordination. In the ordination panel A) each point represents a sample colored according to depth and the shape indicates creek location. B) The results of dispersion tests and Andonis2 PERMANOVA. C) Phylum level composition for each core, with cores collected from Sweeney Creek on the left and West Creek on the right.

### 3. Distinct distributions and depth dependent functional potential among MAGs within network modules

Based on MAG relative abundance, we identified a network containing 1547 positive connections and zero negative connections among 226 of the 377 MAGs (Fig. 4A). The network had a power-law R^2^ of 0.97, indicating a scale-free network, with most MAGs in the network having few connections. Most of MAGs were peripheral nodes in the network, but one MAG (Dehalococcoidales – UBA5760) was identified as a module hub and 15 MAGs were classified as connectors (Fig. S4). Among the connectors were two Bathyarchaeia classified as BA1, two Dehalococcoidia - AB-539-J10, Bacteroidia – vadinHA17, and Planctomycetota – SG8-4 MAGs. Connector nodes represent potential metabolic connections among modules with a widespread distribution, making them interesting targets for metabolic handoffs that sustain multiple interacting groups. Modules 1, 2, and 7 contained the largest number of MAGs with a high degree of connectivity and a diversity of taxa that included 14, 23, and 6 phyla respectively (Fig. 4A). Connectivity between module 4 and 5 was sparse in comparison to the large number of positive connections among modules 1, 2, and 7.

**Figure 4.**
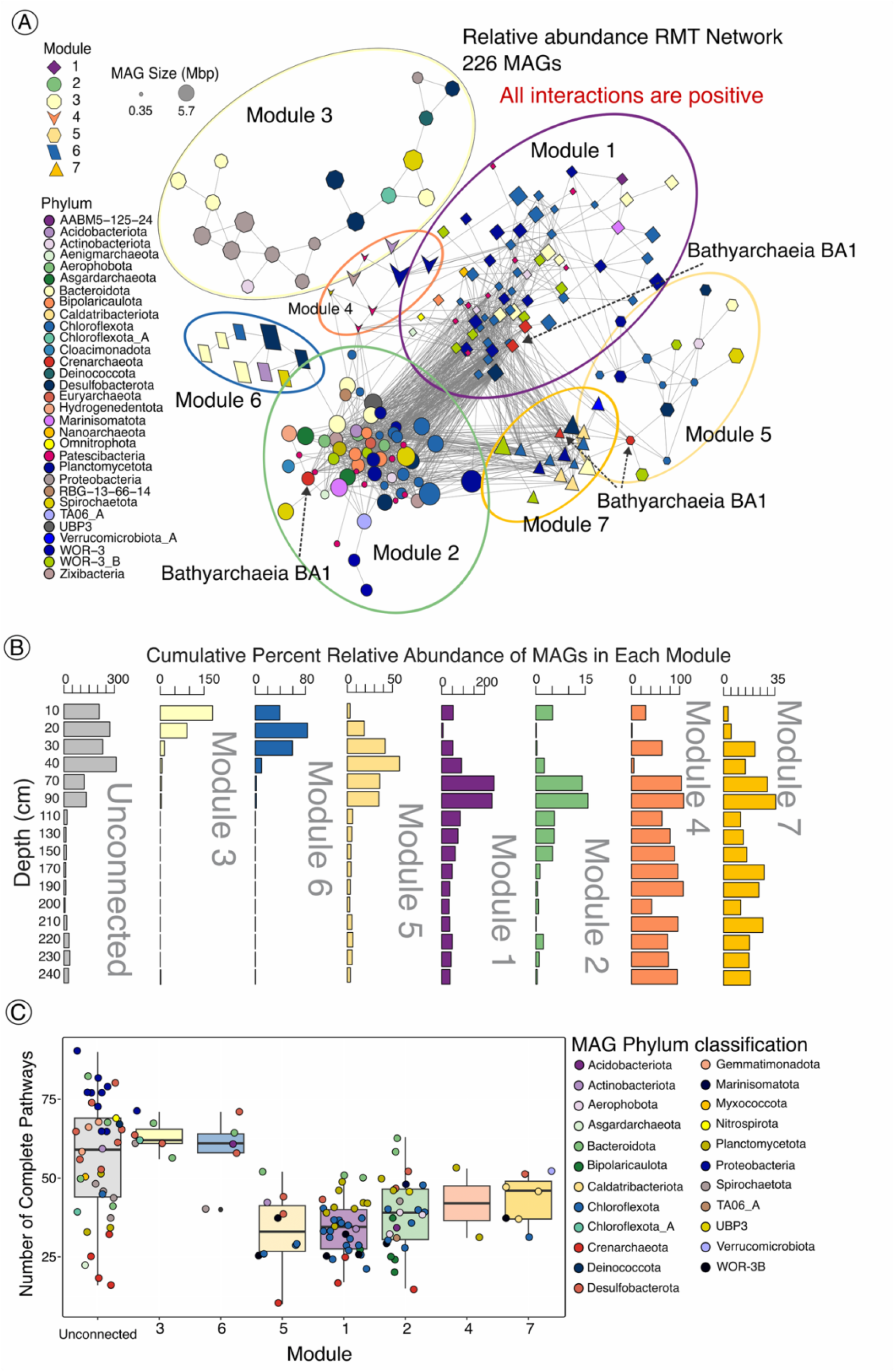
A.) Random matrix theory derived cooccurrence network, based on MAG relative abundance. The point size scales with genome size, the color indicates the phylum classification, and the shape identifies the module. All connections are significant and positive. Ellipses are used to clarify the modules. B.) Cumulative relative abundance of MAGs in network modules with eight or more members. The color of the bars are used to draw attention to the patterns and align with the module ellipses C.) The number of complete pathways plotted for each MAG and box-whisker plots for the median, 25^th^ and 75^th^ percentile lines for each of the modules of interest.

Each collection of MAGs within a network module had a distinct distribution with depth in the sediment (Fig. 4B). The unconnected MAGs with no significant correlations were primarily distributed within the top 40 cm (Fig. 4B). Modules 3 and 6 were also localized above 40 cm while module 5 peaked at approximately 40 cm. However, modules 1,2, 4, and 7 had peak abundances at 70 and 90 cm (Fig. 4B). The depth distribution of MAGs was reflected in the average number of complete metabolic pathways, which were most abundant in the unconnected MAGs and in modules 3 and 6 (Fig. 4C). Collectively, the MAGs in these modules had significantly larger genome sizes and number of complete pathways compared to the modules with a deeper distribution (modules 1, 2, 4, and 7) (Fig. S5). There was no difference in the mean percent completion between MAGs in the shallow and deep modules (Fig. S5).

Pathways for carbon fixation, nitrogen reduction, sulfur oxidation and reduction, chlorate/perchlorate reduction and oxidative phosphorylation were most common in the unconnected MAGs, and modules 3 and 6, typically found in shallow sediments (Fig. 5), and were rarely identified in MAGs that were assigned to the deeper modules. The prevalence of cytochrome oxidases was also more common in MAGs found in shallow sediments (Fig. 5). Iron/manganese and selenate reduction were similar in proportion across all modules along with carbon fixation via the Wood-Ljungdahl pathway, demonstrating the universal importance of these functions within the entire sediment profile examined here (Fig. 5).

**Figure 5.**
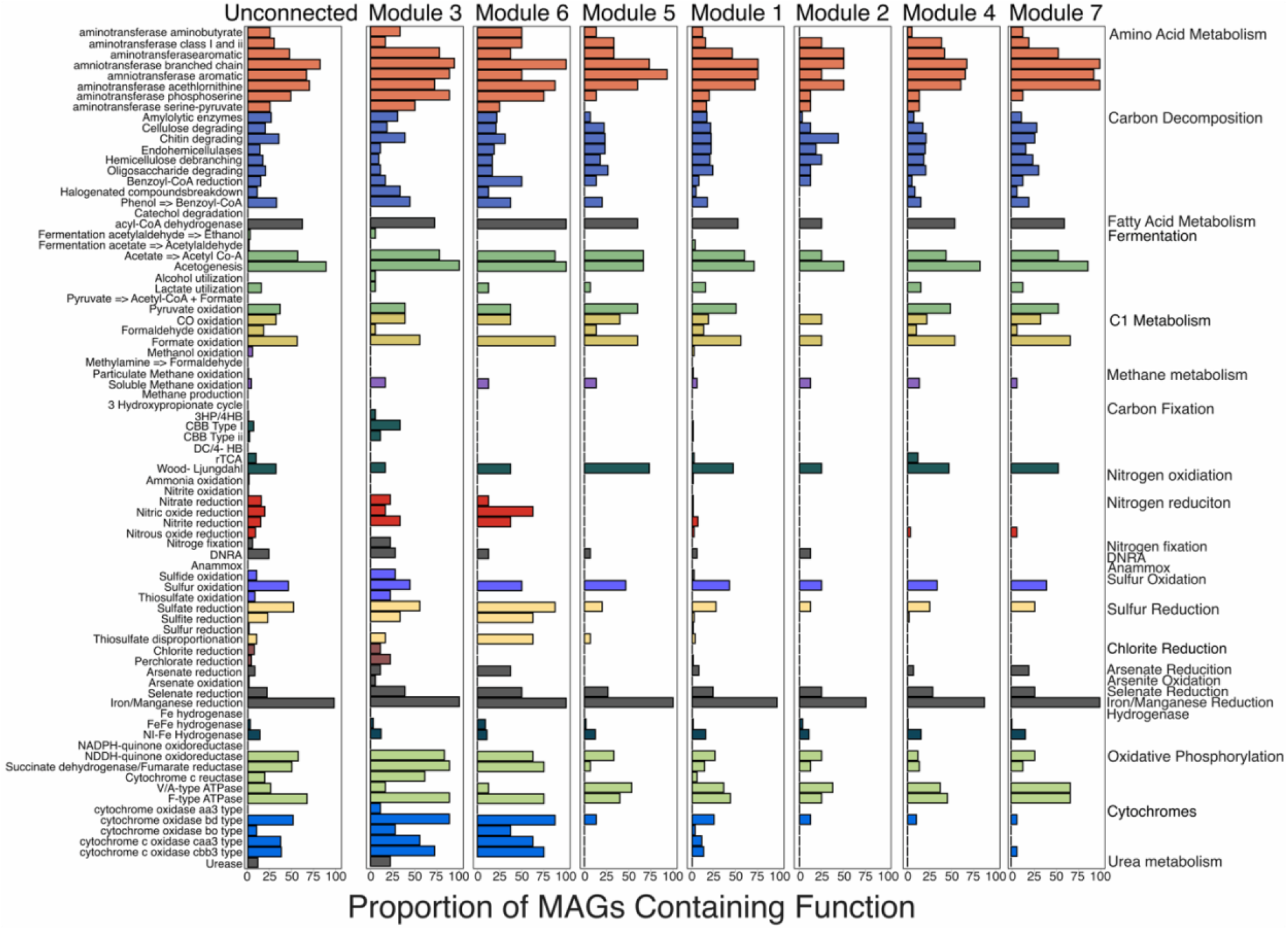
Functions detected within each of the network modules, with individual functions on the left and functional category on the right. The bars are colored according to the functional category. Each of the separate bar plots represents the proportion of MAGs that contain the function within a network module. The modules are listed in the same order as Fig. 4, where the collective relative abundance distribution of MAGs goes from shallower to deeper (left to right).

Carbon decomposition potential was identified in roughly 20% of MAGs in all modules and unconnected MAGs. Specific targets for carbon decomposition included chitin, cellulose, and aromatic compounds through the Benzoyl-CoA pathway (Fig. 5). We found no significant difference in the number of glucoside hydrolase (GH) counts in MAGs among modules, even after correcting for the number of genes in the MAG (p = 0.885). Counts of GH enzymes were highest within the unconnected Planctomycetota MAGs. Six of these MAGs contained over 100 genes with hits to 29-55 unique GH enzymes (Fig. S6A, Table S2). The Planctomycetota within modules 1, 2, 4, and 7 also contained over 100 GH counts from a similar diversity of CAZymes (Table S2). Additional MAGs containing large numbers of CAZymes within the unconnected and shallow modules (3, and 6) included Calditrichota, Bacteroidota, and Spirochaetota (Fig. S6A). CAZymes classified as N-acetylglucosaminidase (NAG) extracellular enzymes, were among the most commonly identified (Table S2). Additional abundant enzymes detected across our collection of MAGs included GH13 involved in starch and sucrose metabolism, GH2, which targets a diverse group of carbohydrates, and GH23, which targets β-1,4-glycosidic bonds in bacterial cell walls. Peptidase homolog counts were significantly different among the MAGs in the shallow (unconnected and modules 3 and 6) and deeper modules (Fig, S6B, Table S3). Proteobacteria, Desulfobacterota, and Bacteroidota in the unconnected and module 3 MAGs contained the highest counts. The most abundant peptidases were C26 which can degrade peptidoglycan found in bacterial cell walls, M38 involved in protein repair and degradation, S33 a prolyl aminopeptidase and I87 and S14 which modify peptides and regulate specific protein degradation respectively (Table S4).

### 4. Bathyarchaeia BA1 subnetwork connected MAGs are widespread and play an active role in decomposition of complex carbon and aromatics

#### 4.1 Consistent relative abundance among members of the Bathyarchaeia BA1 subnetwork

The potentially important ecological role of Bathyarchaeia BA1 MAGs, indicated by their role as connectors in the network (Fig. S4) and the global significance of this family to carbon cycling (Hou et al. 2023) made it an intriguing target to study in greater detail. If connectors are important to the stability of the community, Bathyarchaeia BA1 metabolism may be important to the survival of other MAGs through metabolic handoffs. Additionally, environmental filtering could have led to their prevalence across the entire sediment profile leading to a positive correlation across multiple modules. To tease apart these details, we focused on the Bathyarchaeia BA1 MAGs and their connections to other MAGs in the network to examine the functions that would support survival within the energy limited subsurface. We created a subnetwork of MAGs that were directly connected to the Bathyarchaeia BA1 MAGs, resulting in a total of 33 MAGs (Fig. S7A). The Bathyarchaeia BA1 subnetwork MAGs represented 20-30% of all mapped reads in the samples below 150 cm in both the metagenome and metatranscriptome and the MAGs were compositionally similar across samples (Fig. 6). The subnetwork captured 10 out of the 11 connector MAGs (Table S1), including taxa with a well-documented presence in energy limited anaerobic environments; Dehalococcoida - AB- 539-J10, Dehalococcoidia – E44-bin88, Desulfobacterota – B25-G16 and WOR3B – UBA3072 (Mara et al. 2023; Shaw et al. 2025). Individual MAGs, classified as connectors, were also among the most abundant (Fig. 6, Table S1). A Desulfobacterota – B25-G16 MAG was the most abundant in the transcriptome and was common and abundant in the metagenome (Fig. 6).

**Figure 6.**
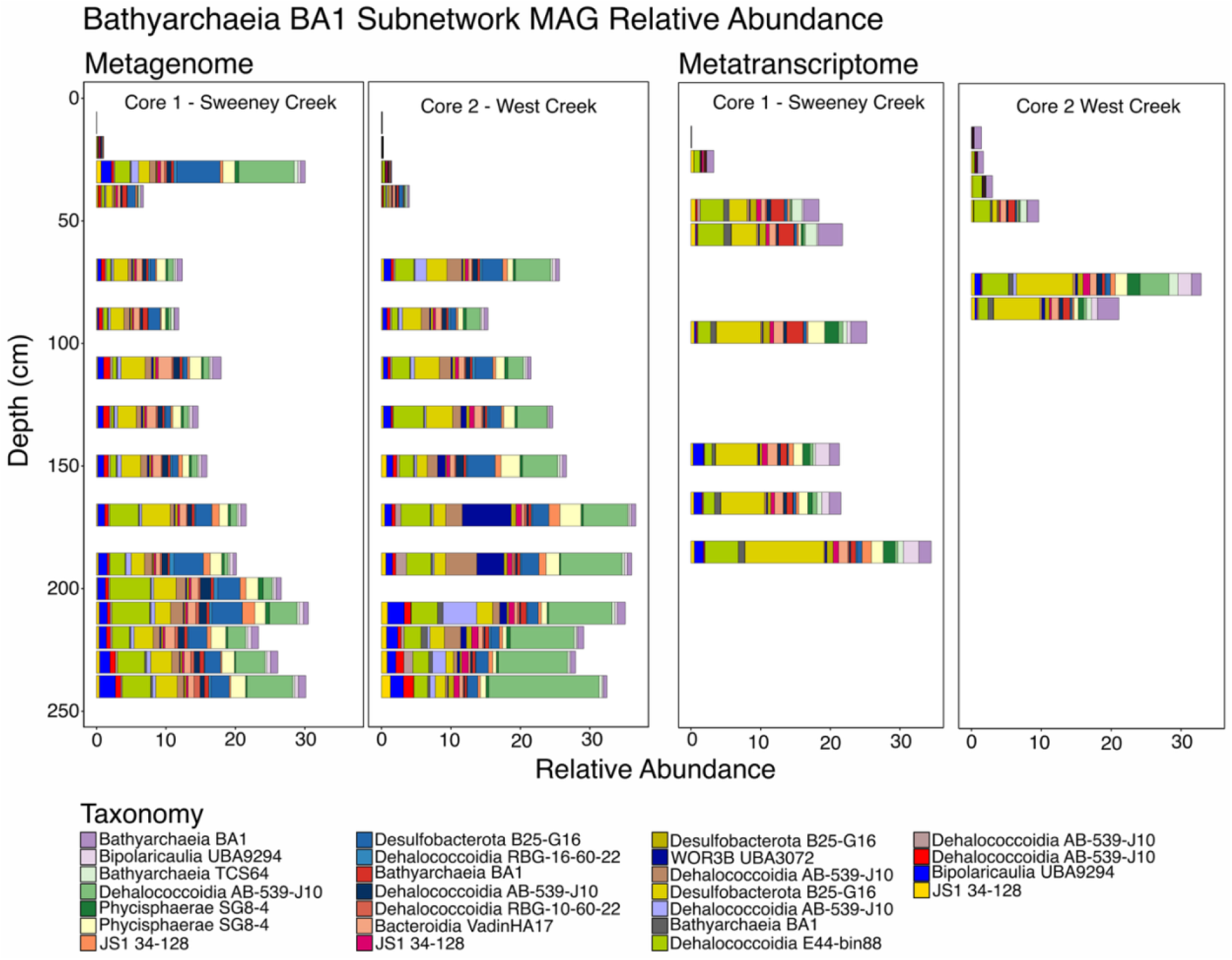
The percent relative abundance of MAGs in the Bathyarchaeia BA1 subnetwork community relative to all mapped reads in the metagenome and metatranscriptome samples in each of the two deepest cores. Only MAGs identified as abundant (mean %relative abundance > 0.1) were shown for the metagenome data and only those MAGs were displayed for the metatranscriptome data. This represents 25 out of the 33 MAGs in the subnetwork.

#### 4.2 Autotrophy, complex carbon degradation, amino acid metabolism, and fermentation were abundant and active in the Bathyarchaeia BA1 subnetwork

We identified several functions among the Bathyarchaeia BA1 subnetwork community that are known to support microbial life under energy limited conditions. Amino acid metabolism, fermentation, acetate oxidation, formate oxidation, complex carbon degradation and the Wood-Ljungdahl pathway contributed the most to cumulative metagenomic metabolic weight of this community (Fig. 7B). Fermentation, complex carbon degradation and CO oxidation were the highest cumulative metabolic weight in the metatranscriptome (Fig.7B). However, Iron oxidation, fermentation and hydrogen generation carried more metabolic weight in the metatranscriptome compared to the metagenomic samples (Fig. 7B). We highlight several abundant MAGs with strong individual contributions to these functions in the metagenomic and metatranscriptomic data as exemplars (Fig. 7A, B) and highlight their distribution across all cores (Fig. 7C).

**Figure 7.**
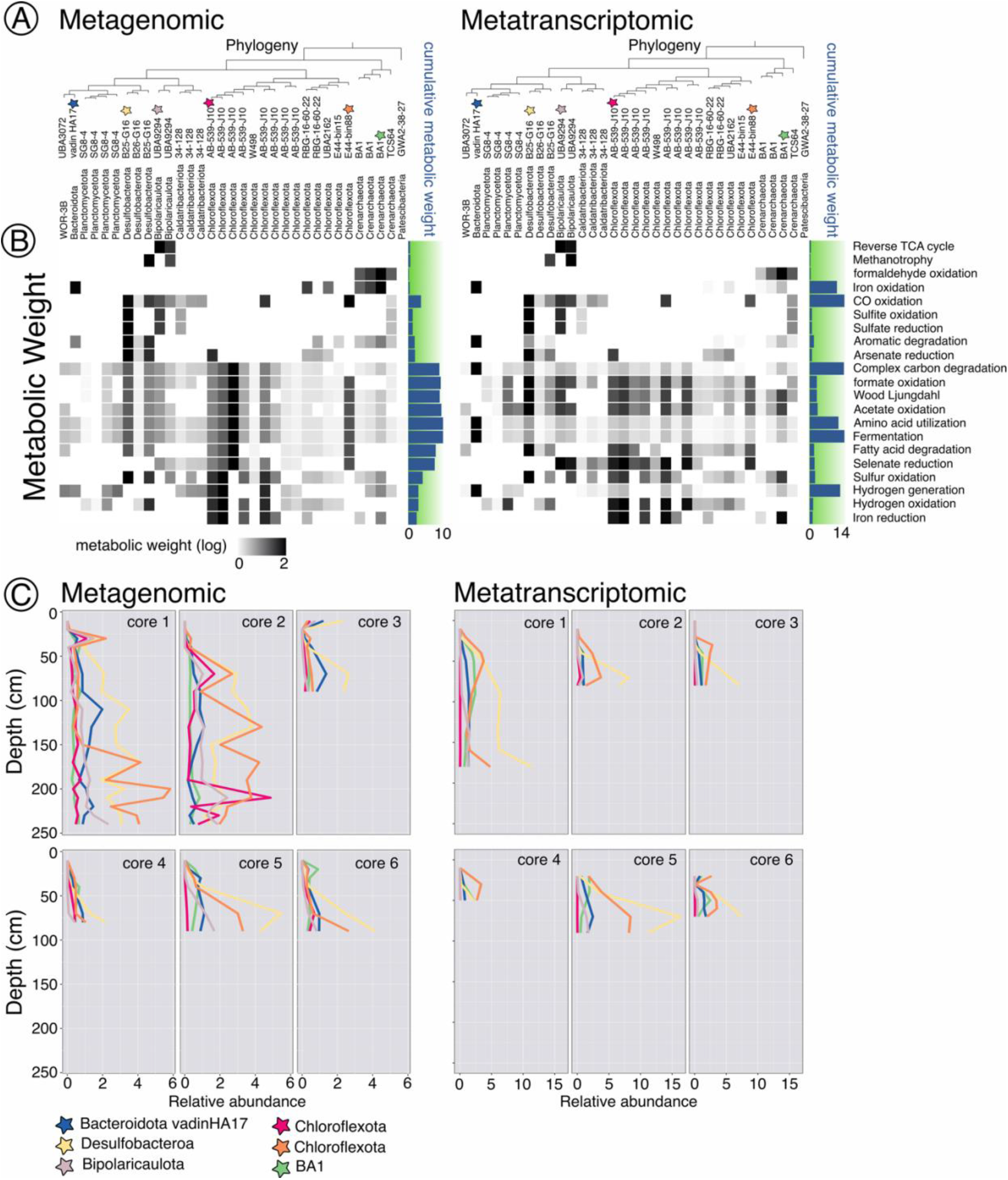
Bathyarchaeia BA1 subnetwork of metagenomic (MG) and metatranscriptomic (MT) functions. A.) The phylogenetic relationship among MAGs according to single copy core genes with family and phylum level classification. Stars are used to highlight several MAGs that have high metabolic weight in the MG and MT data. B.) Heat map for the metabolic weight contribution in the MG and MT to each of the functional categories displayed on the right-hand side of the figure. The blue bars represent the cumulative metabolic weight of the functional category within either the MG or MT data. C.) The relative abundance of the starred MAGs from panel A. Corresponding cores are shown for each dataset and labeled within each individual panel. Relative abundance in this case is the percentage of reads mapped to that MAG relative to all mapped reads.

Genes for heterotrophic and/or autotrophic acetogenesis (homoacetogensis), through the Wood-Ljungdahl (WL) pathway, were detected in 25 of the 33 genomes in the Bathyarchaeia BA1 subnetwork. (Fig. S7C). Within the subnetwork, the metabolic weight of pathways involved in acetate metabolism including WL, formate oxidation/reduction, and acetate oxidation/reduction were similar in cumulative metabolic weight (Fig. 7B). METABOLIC scored a Dehalococcoidia AB-539-J10 MAG as having the highest contributing metabolic weight for all three pathways in the metagenomic data. A Desulfobacterota B25-G16 was the largest contributor to these pathways in the metatranscriptomes (Fig. 7B) and was among the most abundant and active MAGs in the subnetwork (Fig. 7C).

Individual genes associated with acetogenesis identified among the Bathyarchaeia BA1 subnetwork community were acetyl-CoA dehydrogenase (*acdA*), acetate kinase (*ack*), and phosphate acetyltransferase (*pta*) (Table S5). *acdA* was the most often detected gene indicating acetogenic metabolism, and *acs* (Acetyl-CoA synthetase) was commonly detected within the same MAG (Table S5). CO oxidation made up nearly 14% of the cumulative metabolic weight in the metatranscriptome and potentially couples electrons to the Wood-Ljungdahl (WL) pathway. However, CO oxidation was not observed at the same level of cooccurrence with WL in the same MAG as formate and acetate metabolism (Fig. 7B). Gene annotation of a complete acetate metabolism pathway, the WL pathway, and a formate dehydrogenase gene required to convert CO_2_ into formate within the same MAG provides evidence for acetogenic metabolism.

Complex carbon degradation genes were abundant and active within the BA1 subnetwork community (Fig. 7B). Bacteroidia– vadinHA17 MAG was the strongest contributor to the cumulative metabolic weight of this functional category (Fig. 7B). The CAZyme profile for this MAG contained 93 gene hits to 36 unique GH genes including eight genes targeting hemicellulose, cellulose, chitin, and alpha amylase (Table S2, S5). The closely related Phycisphaerae SG8-4 MAGs contained similar diverse targets for complex carbon degradation (Table S2, S5). Other MAGs in the community that contributed to the overall metabolic weight of complex carbon degradation targeted a smaller number of carbon compounds that reflects potential specialization on a particular carbon source (Table S5).

Aromatics were a target of the Bacteroidia vadinHA17, especially in the metatranscriptomes, with additional contributions from the Bathyarchaeia BA1 subnetwork, including Desulfobacterota B25-G16. The pathways identified by METABOLIC for aromatic decomposition included metabolism of phenol to benzoyl Co-A and benzoyl Co-A reduction (Table S5). Bacteroidota – vadinHA17 also contributed the most to the cumulative metabolic weight for iron oxidation with additional contribution from the Bathyarchaeia BA1 group (Fig. 7B) in both the metagenomes and metatranscriptomes.

#### 4.3 Functional profiles of the Bathyarchaeia BA1 subnetwork indicate metabolism of complex carbon

M2M identifies added value metabolites that are produced through complementary metabolisms among the microbes in a minimal community. A minimal community represents a group of MAGs required to produce the entire set of metabolites in a MAG collection. We found 82 metabolites that were producible by each of the 33 MAGs in the subnetwork (core functions) and 1098 total metabolites that could be produced by the collection of organisms in the subnetwork (collective functions). The added value of cooperation within a minimal community required 22 of the total 33 MAGs to produce an additional 228 compounds. These 228 compounds were only possible when all members of a minimal community had the potential to exchange metabolites or conduct sequential utilization of substrates. Redundant members of the Bathyarchaeia BA1 subnetwork that were identified in at least one, but not all minimal communities included Dehalococcoidia – RBG-16-60-22, JS1 34-128, Dehalococcoidia AB-539-J10, and Dehalococcoidia UBA5760.

Among the products of cooperative metabolism were aromatic compounds, related to styrene, toluene, phenylacetate and betaine degradation (Table S6). One of the four Benzoyl-CoA reductase (*bcrABCD*) subunits that are required for ring cleavage of aromatics was identified in the genomes of three Desulfobacterota - B25-G16, a novel order of Dehalococcoidia – E44-bin15, and Bathyarchaeia - BA1 and the composition of the genes were complementary (Table S7C). None of the MAGs had the complete set of Benzoyl-CoA reductase genes. Genes to produce benzoyl-CoA from phenol (*ubiX*/*bsdC*) occurred in two of the Desulfobacterota – B25-G16, and all Bathyarchaeia and Bacteroidia vadinHA17 MAGs (Fig. S7C). Phenylacetyl-CoA, which uses a ring cleavage system distinctly different from Benzoyl-CoA was also identified as a metabolite putatively produced through functional complementarity within the subnetwork (Table S6).

#### 4.3 SNVs are structured by depth and location among MAGs in the Bathyarchaeia – BA1 subnetwork community

We analyzed the composition of sample specific nucleotide variability within each of the MAGs in the Bathyarchaeia BA1 subnetwork community. Six of the MAGs had sufficient coverage throughout the depth profile to test for the effect of depth and site on SNVs frequency (Fig. S8). In several MAGs, the overall variability declined with depth, but this trend was not observed for all MAGs (Fig. 8A-F). SNV frequencies were generally not correlated with depth alone (Fig. 8, Table S8) and even when depth was significant, the R^2^ was very small (Table S8). Site, however, was a significant variable explaining the observed differences in SNV frequency among samples. The interaction between site and depth was significant and the term improved the model for all MAGs, explaining more than 25% of the variability in SNV frequency (Table S8). Visualization of the ordination confirms a loose structuring according to depth and well-defined differences between cores (Fig. 8).

**Figure 8.**
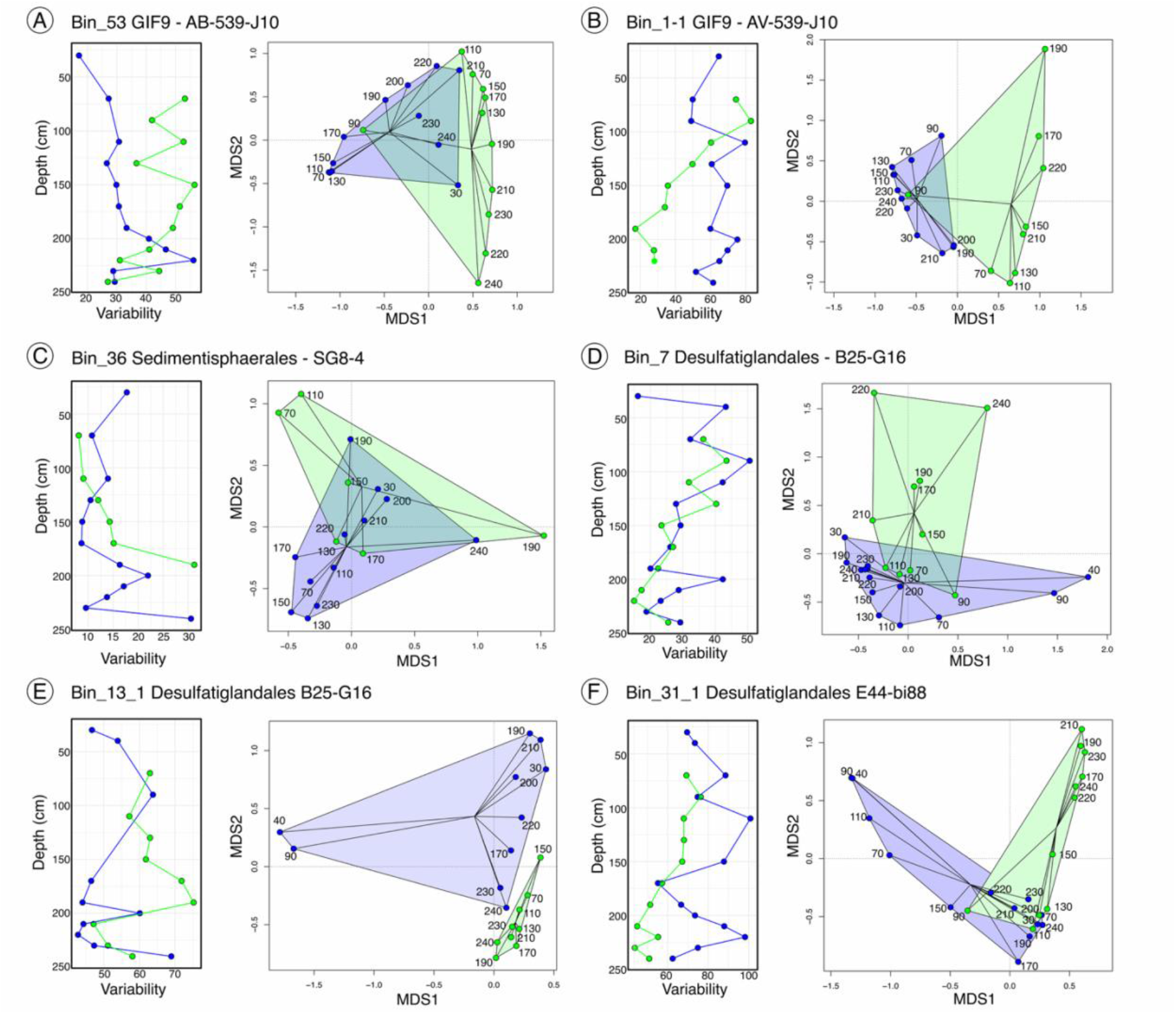
Variability ((number of variants/genome length)*1000) and sample specific variant clustering of six MAGs from the subsurface core community (A-F). The analysis was limited to samples where the mean coverage was > 15X. Each panel contains a plot of the variability with depth. Separate lines are drawn for each of the two cores (blue = Sweeney Creek, green = West Creek). Ordination based on SNV variant frequency at 5000 randomly chosen sites in the genome. Geometric shapes and lines to the centroid are shown for the two cores and color coded according to creek. Statistics for the ordination are located in Table S8.

## Discussion

Microbial communities contribute to the rapid depletion of terminal electron acceptors and biologically available carbon from the surface of salt marsh sediments (Benner et al. 1984; Fogel et al. 1989; Sutton-Grier et al. 2011). The sequential decomposition of carbon in the surface leads to the accumulation of less biologically available compounds for microbial growth within deep sediments (Leadbeater et al. 2021; Fogel et al. 1989). We know relatively little about the microbial communities that inhabit these low energy deep sediments, and the composition and metabolic strategies used by these microbes represents a significant knowledge gap in our understanding of carbon cycling within this blue carbon system. Our study of deep sediment cores addresses several aspects of the ecology and evolution of these communities with a special focus on their contribution to carbon cycling.

### 1. Microbial communities are structured according to depth with a steep transition between 40-70 cm

Studies of microbial communities below 20 – 40 cm in the salt marsh are rare and often employ 16S rRNA amplicon surveys, which limits our ability to identify the boundaries of microbes and their metabolisms. Previous studies of microbial turnover within salt marshes described shallow sediment seasonal and transect compositional changes (Tebbe et al. 2022; Kearns et al. 2019; Yao et al. 2019) or succession along a chronosequence (Dini-Andreote et al. 2014). The composition and stability of microbial communities within salt marsh sediments remains relatively uncharacterized at the depths examined here. Our results from the beta diversity analysis, MAG dereplication, and network analysis all support our hypothesis that the microbial community was significantly shaped by depth. The largest changes in composition were observed in the transition between 40-70 cm in each of the six cores indicating a distinct change in the physical and biogeochemical characteristics of the sediment in this high marsh platform. Most of the MAGs located at and above 40 cm were reconstructed from a single sample and were not correlated with other MAGs, indicating a patchy distribution. Microbial communities from these depths are known to contain high phylogenetic diversity (Bowen et al. 2012) and changes in composition can occur across small spatial scales (Marlow et al. 2021). Depths below 40 cm were characterized by similar community composition and high relative abundance of a relatively small number of MAGs which resulted in our ability to reconstruct their genomes from multiple samples and detect cooccurrence. Many of the MAGs in the subsurface were also detectable in samples above 40 cm, potentially pointing to an origin from surface sediments. These results indicate a relatively low diversity environment below 40 cm that is restricted to a subset of microbes that are also capable of inhabiting niches within shallower sediments (Starnawski et al. 2017).

The distinct change in community composition between 40-70 cm could be explained by several physical and biogeochemical characteristics of the sediment. Plant roots and rhizomes along with bioturbating organisms can increase permeability in the upper layers of the sediment (Guimond and Tamborski 2021; Moffett et al. 2012; Xiao et al. 2019) allowing for movement of solutes. In salt marshes, hydrological effects are strongest in the upper 100 cm and 10-15 m from the creek bank (Nuttle and Hemand 1988) and low porosity, small particle size, and low circulation can restrict the movement of water in deeper sediments (Guimond and Tamborski 2021). Additional variability in the hydrology of the marsh, primarily at the surface, results from flushing of the interior during spring tidal flooding, advection, and the seasonal effects of plant evapotranspiration on water movement (Hussey and Odum 1992). The resulting movement of water carrying terminal electron acceptors, carbon sources, and fluid associated microbes through pores in the sediment can result in rapidly changing conditions favoring habitat generalists in the surface (Chen et al. 2022) and leading to decomposition of otherwise stable pools of carbon in the upper 40 cm (Bulseco et al. 2019). The microbial community shifts observed in our study point to a steep gradient in salt marsh sediments from one shaped by dynamic hydrology and plant driven interactions near the surface, to a more stable environment that persisted for more than two meters below the surface.

### 2. Cooccurrence, genome size, and reduced metabolic potential suggests subsurface communities are shaped by environmental filtering

Our study of the vertical sediment profile recovered several pieces of evidence that suggest environmental filtering is likely the underlying mechanism structuring the subnetwork community. First, we recovered MAGs that we assembled independently from multiple depths in the same core that dereplicated into a single MAG. For example, the Bacteroidota vadinHA17 was recovered independently from 15 samples and the candidate phylum TA06 had a single population that was recovered from 24 of the 55 samples, including every depth fraction except the shallowest. For dereplication to occur at this scale (> 95% identity among MAGs in the dereplicated cluster), genome populations would have experienced strong environmental selection followed by limited growth and/or mutation rates and continued selection over the nearly 3000 years of separation that exists between the surface and bottom of the deepest cores. Second, we observed a consistent correlated pattern among MAGs in sediments below 40 cm, reflecting stable conditions at these depths.

The presence of unique modules within the ecological network could be indicative of distinct ecological niches within the community (Xie et al. 2024). Our study identified 7 modules and the MAGs within these modules shared a unique distribution in the environment that could reflect adaptation to a niche present within a specific depth. Two of the modules contained MAGs that were primarily detected at or above 40 cm. Additionally, the unconnected MAGs were generally more abundant within the upper 40 cm and MAGs with a 40-70 cm distribution in the sediment. A large proportion of MAGs in these groups contained striking genomic versatility including the capacity to utilize multiple terminal electron acceptors for oxidation of diverse carbon sources, and conduct fermentation and carbon fixation (Table S9), metabolisms that are commonly associated with salt marsh microbial communities. Many of the MAGs with a 40-70 cm distribution contain the genomic potential for each of these metabolic pathways within a single organism, as observed previously (Vineis et al. 2023). This metabolic flexibility allows each of the members of the community to inhabit a variety of conditions, consistent with the dynamic nature of these sediments. Although we did not evaluate the presence of available nitrate, sulfate, oxygen or other potential electron acceptors in the profile, other studies have demonstrated that they are rapidly consumed, and often undetectable below the first few centimeters of the surface (Brodersen et al. 2019; Sørensen et al. 1979). Our study indicates that many of the same microbial populations dependent on carbon and nutrients from the upper layers are able to persist, in many cases, up to at least 40 cm.

All other network modules contained MAGs that were primarily abundant below 40 cm. The metabolic potential of the MAGs reflects the conditions beyond 70 cm, which are likely to contain relatively low-quality carbon sources and low concentrations of terminal electron acceptors. The dominant taxa at these depths were Planctomycetota, Chloroflexota, Desulfobacterota, and Bacteroidota, which are known to inhabit energy poor systems (Xamxidin et al. 2025; Rezaei Somee et al. 2026). MAGs with deeper distribution had reduced genome sizes and reduced functional potential despite high completion estimates and genome coverage when compared to MAGs with a peak distribution in the upper sediment layers. These results support our hypothesis that deeper sediment communities are functionally streamlined due to environmental filtering.

Abundant MAGs below 40 cm maintained localized distributions and formed statistically significant correlations due to the stability of their occurrence patterns across each of the cores. This could indicate localized conditions that cooccurring members are adapted to, but there were no obvious metabolic pathways or taxa associated with the individual modules. It is possible that occasional advection could lead to restructuring of the subsurface community (Zhang et al. 2022), but they could also reflect the conditions at the surface prior to burial followed by environmental filtering, which has been observed in other marine sediments (Marshall et al. 2019). The underlying mechanism that led to the formation of distinct relative abundance peaks among cooccurring members remains an intriguing outstanding question.

We could not address our hypothesis aimed at testing the differences in connectivity of MAGs between shallow and deeper sediments because of the low success of MAG reconstruction from surface sediments. However, this is likely due to the high diversity within these sediments (Bowen et al. 2012). This does not negate the potential for metabolic handoffs and important interactions within the surface but simply indicates they are more difficult to tease apart using metagenomic sequencing and MAG reconstruction alone. It is possible that additional assembly and binning approaches, including co-assembly, could yield additional MAGs from the upper 40 cm and this approach is deserving of an independent study.

### 3. The Bathyarchaeia BA1 subnetwork MAGs are widespread and active in the sediment indicating an important role in orchestrating complex carbon cycling

Identification of a subnetwork of cooccurring microbes in the sediment sheds light on the ecological and evolutionary forces that shape salt marsh sediment communities. Our results support the presence of a subnetwork that is widespread and abundant in the subsurface. The subnetwork was recovered independently from multiple samples across two different sites, was present in both the metagenomes and metatranscriptomes of the samples, and the M2M analysis indicated that most members of the subnetwork were required to produce the collective metabolites. These subnetwork taxa have been identified in other subsurface communities, demonstrating their broad importance beyond salt marsh sediments. Bathyarchaeia are widespread in subsurface communities where they play a diverse role in carbon cycling (Hou et al. 2025), the Chloroflexota genus AB-539-J10 was similarly widespread within the deeper clay layers of coastal sediments (Sun et al. 2025). Bacteroidota family vadinHA17 is an important member of acetate producing consortia (Yin et al. 2022), and Bipolaricaulota (Acetothermia) family UBA9294 are deeply branching ancient members of acetogenic anaerobic bacteria (Colman et al. 2022).

The metabolic capacity for carbon cycling among the Bathyarchaeia BA1 subnetwork supports our assertion that the sediment below 40-70 cm is a nutrient poor system where metabolic complementarity enables sequential decomposition of complex carbon. The gene expression data indicating these MAGs in this network are active, provides support for continual cycling of organic matter in these sediments, supported by one of the most ancient carbon fixation pathways developed by life on Earth, the WL pathway. Syntrophic interactions between a bacterial fermenter and CO_2_ reduction are often cited as important to the decomposition of aromatic carbon (Dai et al. 2024; Qiu et al. 2008). The collective functional profile of the Bathyarchaeia BA1 subnetwork suggests that the decomposition of complex organic carbon is conducted by members of the Planctomycetota, WOR3, JS1 and others, which is complemented by the decomposition of aromatics by Bathyarchaeia, Deslufatiglandales, Bacteroidales and Chloroflexota. Benzoyl-CoA is a known intermediate involved in the anaerobic decomposition of aromatic compounds toluene, phenol, ethylbenzene and benzoate (Carmona et al. 2009; Porter and Young 2014). Under anaerobic conditions, reduction of benzoyl-CoA leads to the production of acetyl-CoA which can enter central metabolic pathways for growth and energy conservation (Porter and Young 2014).

In many oligotrophic sediments, the products of secondary fermentation, including H_2_ and formate, are used by methanogens. However, our data suggest that in this system these products are consumed by a diverse group of acetogens. Acetogens grow on organic materials derived from cellulose, lignin, alcohols, hemicellulose, and chitin within energy depleted sediments (Lever 2012; Yu et al. 2018), and there is evidence for the use of lignin as an energy source in Bathyarchaeota (Yu et al. 2018). It is possible that methanogens do exist within the sediments here and we failed to reconstruct them. Additional analysis and experiments will be required to identify the potential importance of methanogenesis in these sediments. However, identification of the abundant and active members of the Bathyarchaeia BA1 subnetwork sheds light on carbon cycling pathways employed by microbial communities in this system.

### 4. Active microbial populations are shaped by depth and origin in the estuary

The depth of sequencing and occurrence of MAGs across many of the deep core samples allowed us to conduct an analysis of SNVs, which is rarely possible due to the generally low coverage of individual genomes in salt marsh sediment metagenomes (Graves et al. 2016; Vineis et al. 2023; Jones et al. 2019). Our analysis provides insight into whether populations within the deep salt marsh sediments analyzed here undergo adaptive evolution and if they accumulate mutations. Our results suggest that the MAGs examined here do not undergo adaptive evolution within the deeper sediments because, according to our dereplication procedure, the same MAGs were recovered from both surface and deep sediments. This could occur if advection or restructuring of the sediment allowed for vertical movement of microbial communities, resulting in a homogenized population throughout the sediment. However, we don’t see evidence for this in our SNV analysis. Instead, we see that the SNV profile of deeper sediments is more similar to each other than to shallow sediment samples, suggesting that there is little vertical movement of these MAGs. Other studies of deep marine sediments have observed similar depth structuring in low energy systems (Walsh et al. 2016; Kirkpatrick et al. 2019; Petro et al. 2019).

## Conclusion

Our study of deep sediment cores within a *Spartina patens* salt marsh identified depth dependent structure of microbial communities and populations. Environmental filtering of microbes was observed with depth, especially below 40 cm in these sediments. Members of the Bathyarchaeia BA1 subnetwork complement each other in their metabolic capacity to enable continued decomposition of organic carbon under energetically limited conditions. The syntrophic interactions identified for the decomposition of aromatics requires further validation, but our results shed light on the potential complimentary pathways and community members relevant to this process. Validating and quantifying the decomposition of complex carbon into microbial biomass represent the next steps towards teasing apart the complex interactions in salt marsh sediments. Identifying these interactions could have broader implications for understanding microbial metabolism in other low energy systems.

## Data Availability

The data described here are described in the U.S. Department of Energy (DOE) Facilities Integrating Collaborations for User Science (FICUS) grant titled “Combining high resolution organic matter characterization and microbial meta-omics to assess the effects of nutrient loading on salt marsh carbon sequestration” (proposal ID = 503576, https://genome.jgi.doe.gov/portal/Comhiguestration/Comhiguestration.info.html). Supplemental data can be accessed using this link to a publicly available github page https://github.com/jvineis/DEEP-CORE-MANUSCRIPT.git.

## Acknowledgements

Samples were collected from the Plum Island Ecosystems LTER, which is supported by NSF (OCE 2224608) and from the NSF funded TIDE project (DEB 1902712). The work (proposal: 503576, doi:10.46936/10.25585/60012592) conducted by the U.S. Department of Energy Joint Genome Institute (https://ror.org/04xm1d337), a DOE Office of Science User Facility, is supported by the Office of Science of the U.S. Department of Energy operated under Contract No. DE-AC02-05CH1123. Additional funding for this project was provided by an NSF PRFB to ANB (1907285), Simons Foundation (Grant #1247132 to ZGC, and Grant #01248016 to JLB), and Department of Energy (DOE DE-SC0024270).

