## Supplementary figures and images for "Decomposition potential and population structure among active microbial communities in deep salt marsh sediments"

### Figure S1

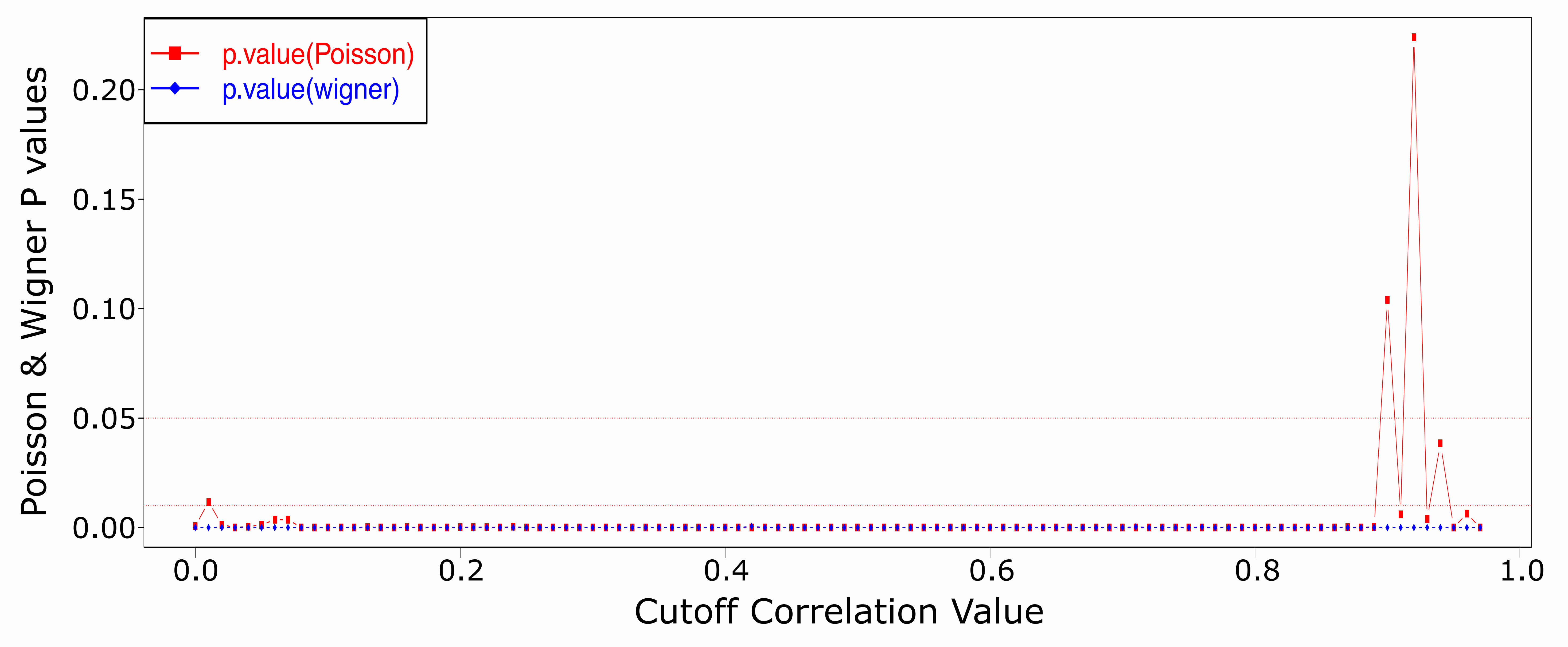

### Figure S2

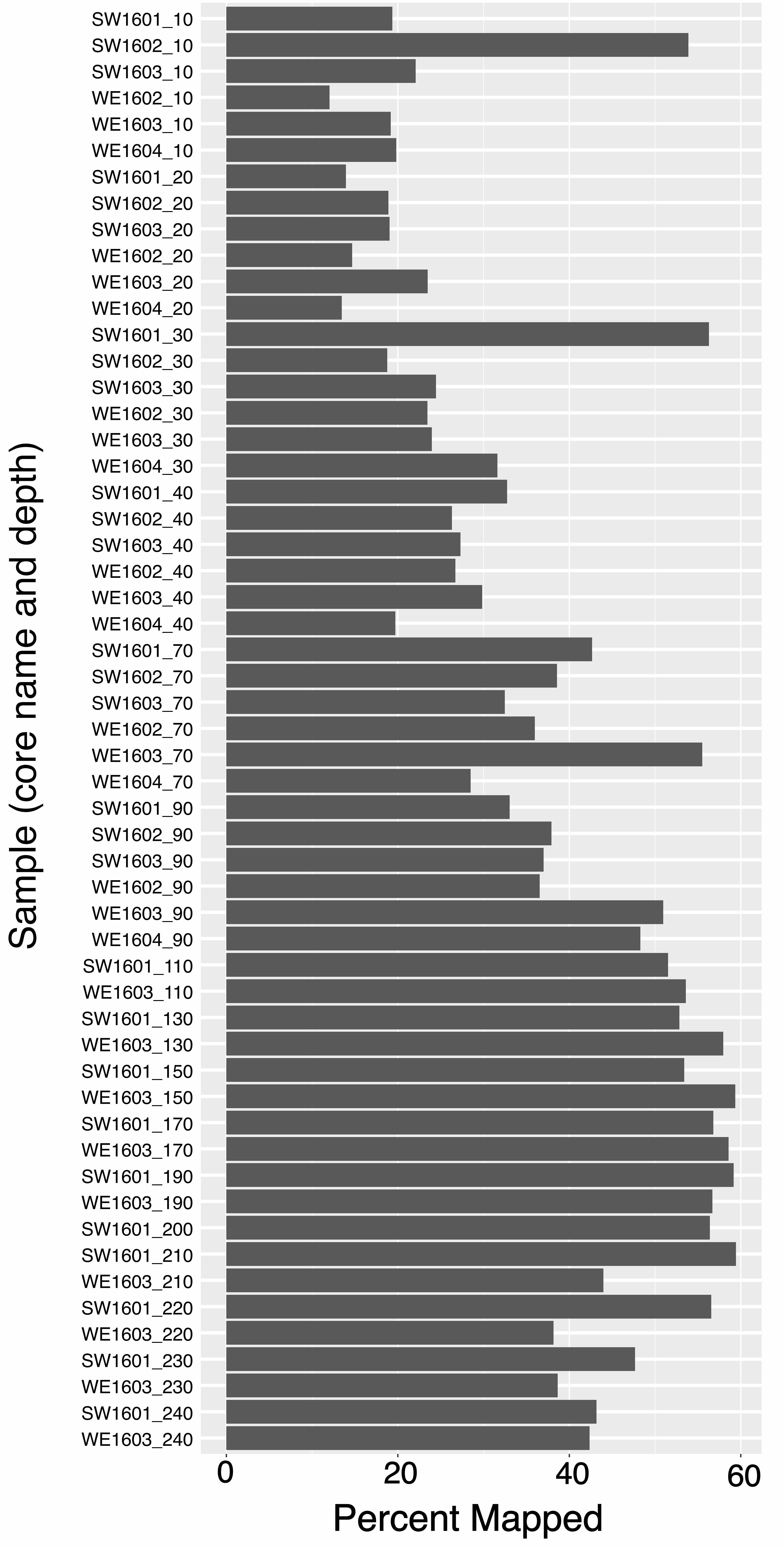

### Figure S3

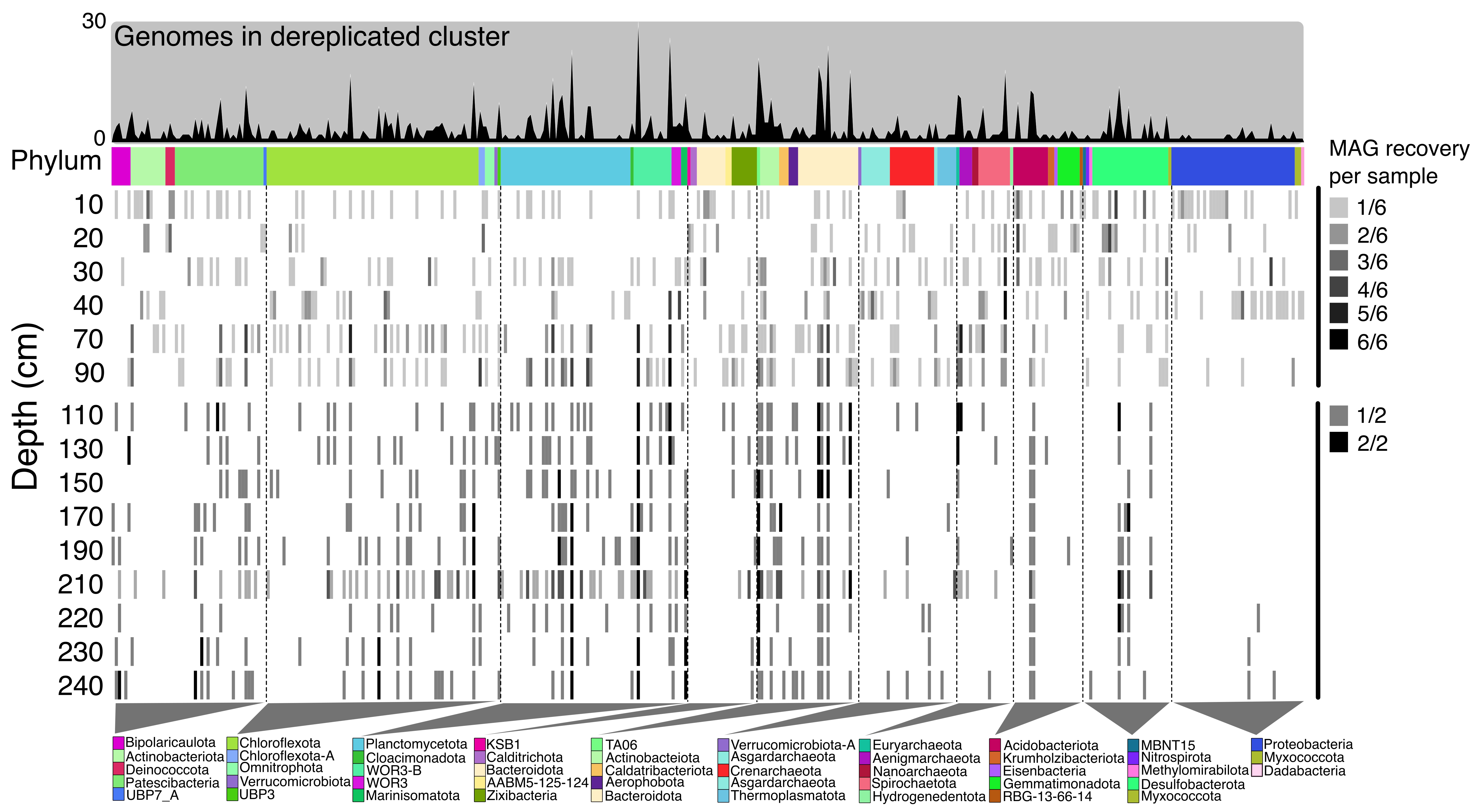

### Figure S4

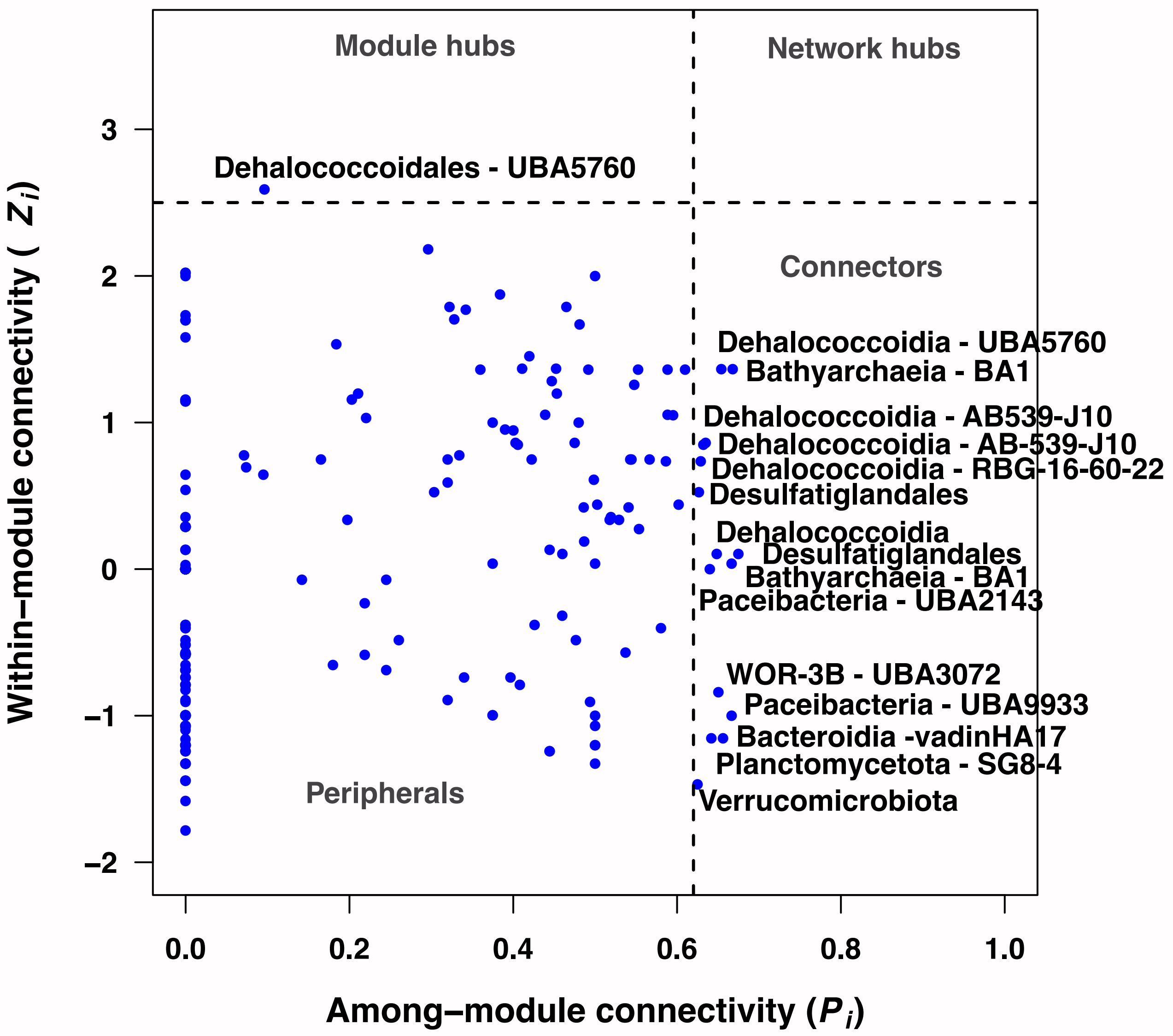

### Figure S5

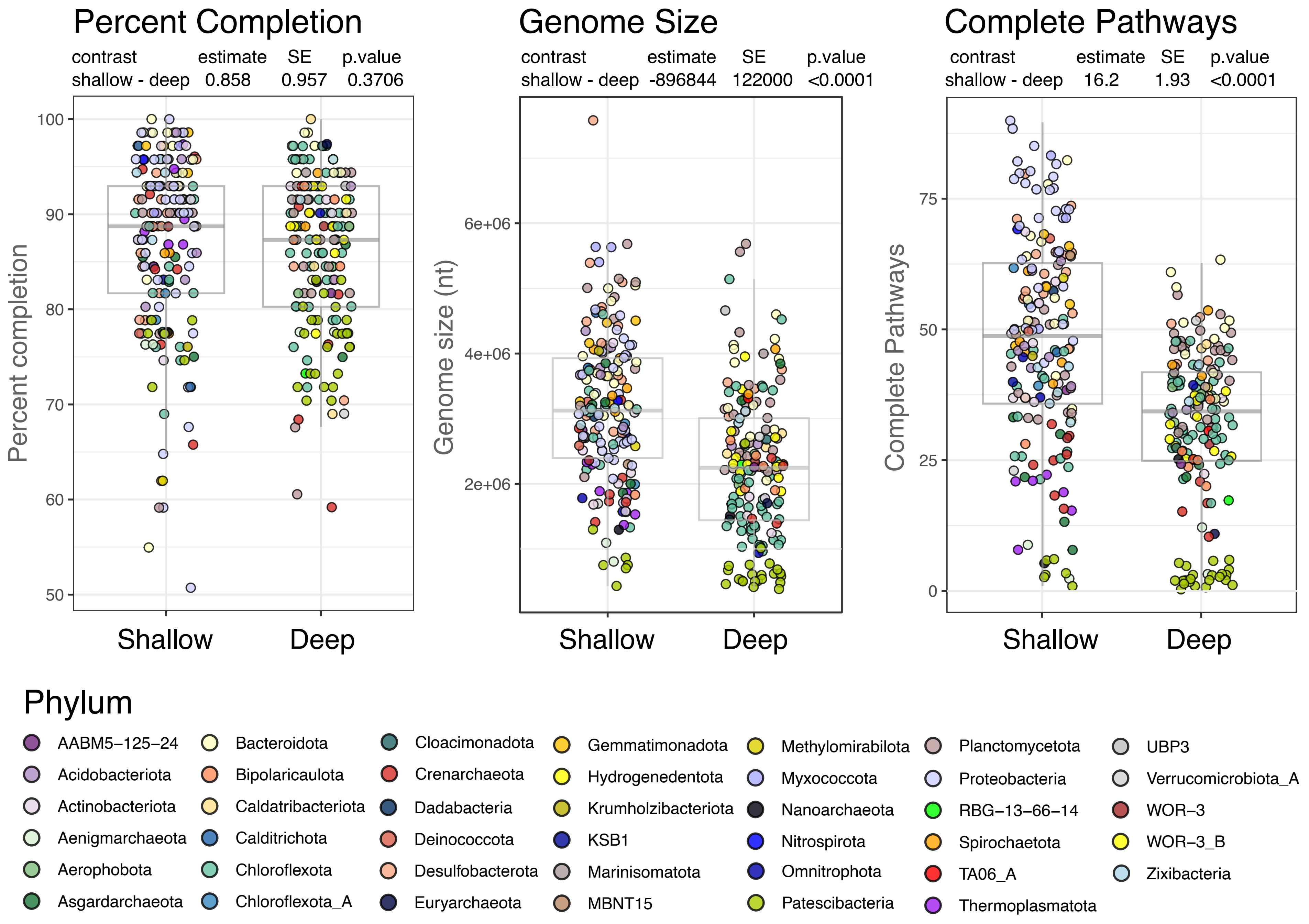

### Figure S6

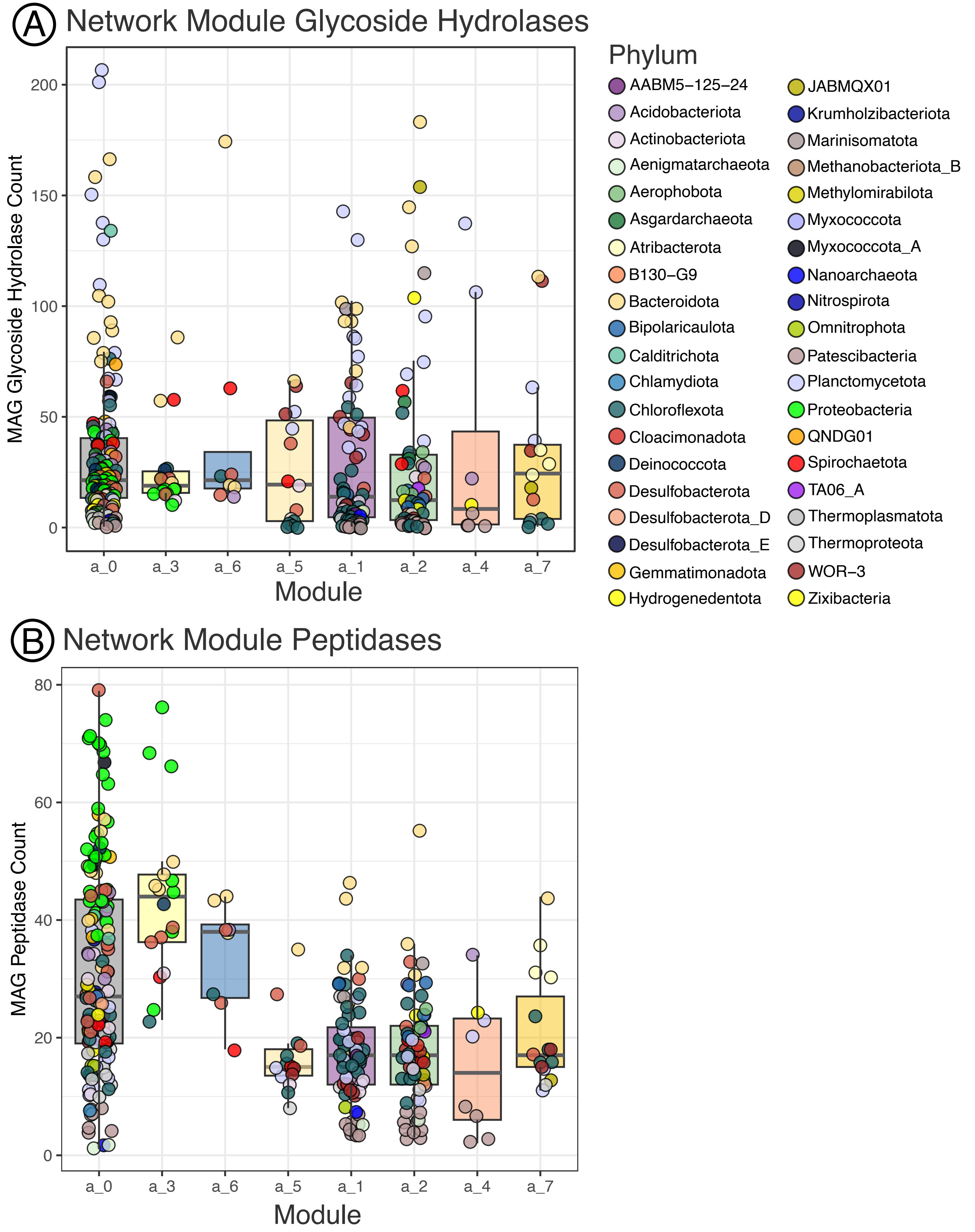

### Figure S7

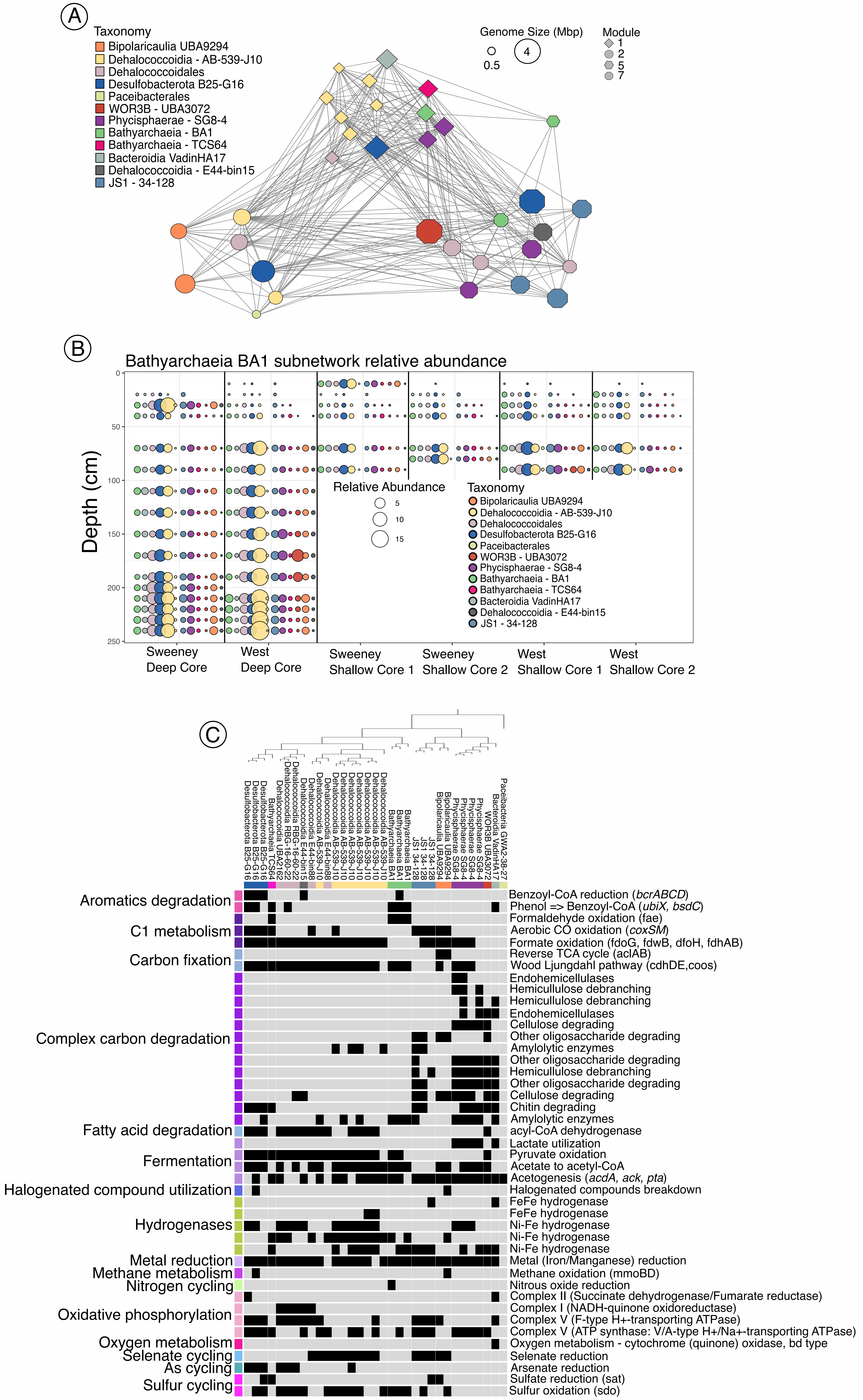

### Figure S8

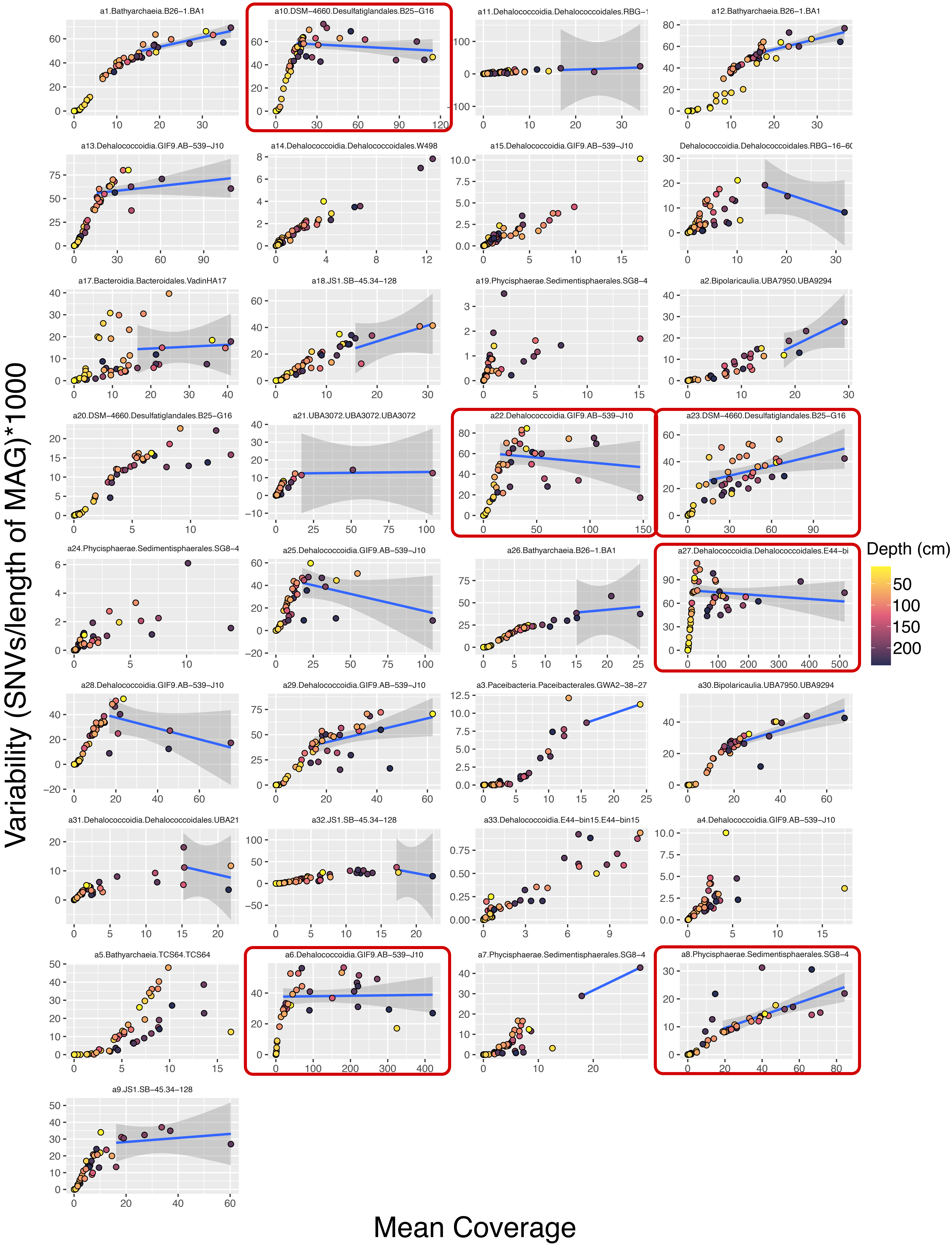
